# Data-driven AI system for learning how to run transcript assemblers

**DOI:** 10.1101/2024.01.25.577290

**Authors:** Yihang Shen, Zhiwen Yan, Carl Kingsford

## Abstract

We introduce AutoTuneX, a data-driven, AI system designed to automatically predict optimal parameters for transcript assemblers — tools for reconstructing expressed transcripts from the reads in a given RNA-seq sample. AutoTuneX is built by learning parameter knowledge from existing RNA-seq samples and transferring this knowledge to unseen samples. On 1588 human RNA-seq samples tested with two transcript assemblers, AutoTuneX predicts parameters that resulted in 98% of samples achieving more accurate transcript assembly compared to using default parameter settings, with some samples experiencing up to a 600% improvement in AUC. AutoTuneX offers a new strategy for automatically optimizing use of sequence analysis tools.

## 1 Main

Biological research increasingly relies on the use of complex software. Many of these tools have tunable parameters that can significantly affect their performance [Majoros and Salzberg, 2004, Frith et al., 2010, DeBlasio and Kececioglu, 2015a, Kovaka et al., 2019, DeBlasio et al., 2020]. A single default setting for these parameters, although carefully tuned by the algorithm designers, is typically not optimal for all inputs of the tools, resulting in the tools’ potential not being fully leveraged. Manually tuning parameters for specific inputs often leads to higher performance, but it is time consuming. A parameter configuration system that can automatically select optimal parameter choices for each input would improve the power of these tools.

Transcript assemblers, software to reconstruct full-length expressed transcripts from the sequenced fragments in a RNA-seq sample, are important bioinformatics tools for RNA-seq analysis that have many tunable parameters. Although a large number of algorithms for transcript assembly have been proposed for assembling different types of RNA-seq samples [Trapnell et al., 2010, Guttman et al., 2010, Tomescu et al., 2013, Pertea et al., 2015, Song et al., 2016, Liu et al., 2016, Shao and Kingsford, 2017, Kovaka et al., 2019, Tung et al., 2019, Liu et al., 2019, Mao et al., 2020, Nip et al., 2020, Zhang et al., 2022a], this task remains challenging due to the diversity of splice variants and biases from the technology such as unevenness of read coverage [Shao and Kingsford, 2017, Voshall and Moriyama, 2018, Zhang et al., 2022a]. Long-read sequencing technologies are able to improve transcript indentification [Cho et al., 2014], however, the challenges of transcript assembly still remain due to the high error rate and sequencing length limits of long-read sequencing [Tung et al., 2019]. Additionally, although previous studies [Kovaka et al., 2019, DeBlasio et al., 2020] have shown that parameter choices are essential to the performance of these algorithms, none of the existing algorithms includes a parameter configuration module to run transcript assemblers with optimal parameter settings.

We introduce AutoTuneX, the first data-driven AI system that automatically recommends parameters for transcript assemblers. For each RNA-seq sample, AutoTuneX predicts the optimal parameters for transcript assembly. AutoTuneX is trained to create a latent embedding of RNA-seq samples, such that samples with similar optimal parameters are positioned closely in the latent space. The training procedure of AutoTuneX has three steps (Fig. 2): (i) selecting a representative sample set, denoted as ℛ_*rep*_, from the RNA-seq sample universeℛ; (ii) determining the best parameter vector for each representative sample 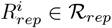 for a specific transcript assembler using Bayesian optimization (BO); and (iii) training the latent embedding through a contrastive learning framework, using information from step (ii) to ensure that representative samples with similar optimal parameter vectors are embedded close together. After the training is complete, the predicted parameters for a new RNA-seq sample are the best parameters of the representative samples that are close to the new sample in the latent space.

We built AutoTuneX separately for two transcript assemblers: Scallop [Shao and Kingsford, 2017] with 18 parameters and StringTie2 [Kovaka et al., 2019] with 8 parameters. A set of values of these parameters is referred to as a parameter vector. We trained AutoTuneX using human RNA-seq samples from Sequence Read Archive [Cochrane et al., 2016], and tested the performance of the predicted parameter vectors on 65 ENCODE RNA-seq samples (ENCODE65), a dataset previously used in Shao and Kingsford [2017]. For each test sample, AutoTuneX generated three sets of predicted parameter vectors of different sizes: top-1 set, top-5 set, and top-30 set. The top-*p* set contains *p* parameter vectors that are the best parameter vectors of the *p* closest representative samples to the test sample in the latent space. We compared these parameter vectors with the default parameter setting for each assembler. When *p >* 1, we assembled the test sample with all parameter vectors in the set and selected the best one, determined by the area under the precision-sensitivity curve (AUC), the same measure used in Shao and Kingsford [2017].

The performance of AutoTuneX is shown in Fig. 1 and Fig. 8. For Scallop, AutoTuneX achieves higher AUC values than the default settings in 53 out of 65 samples for the top-1 set. On average, AutoTuneX delivers a 12% higher AUC than the default settings (Fig. 1c). For the top-5 set, the best AUC of those five parameter vectors surpasses that of the default settings in all samples, with an average AUC improvement of 22%. For the top-30 set, the average AUC is 25% higher than the default settings.

**Figure 1:**
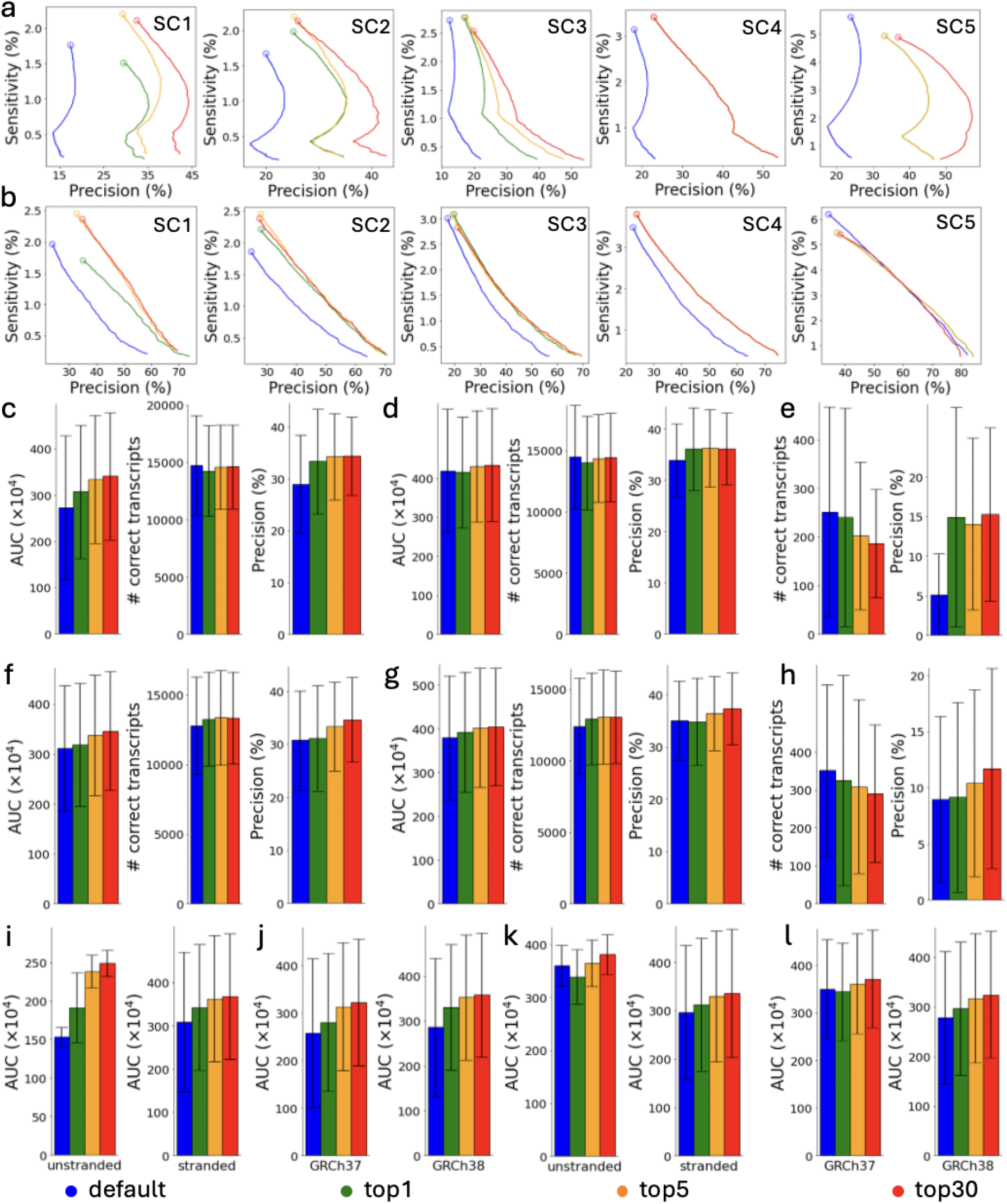
Comparisons between AutoTuneX against the default parameter settings in the top-1, top-5, and top-30 cases for ENCODE65. (a-e,i,j) are results for Scallop, and (f-h,k,l) are results for StringTie2. (a) The precision-sensitivity curves for total (single- and multi-exon) transcripts. The five samples are the samples that achieve the most significant AUC improvement with AutoTuneX for Scallop. The circles represent the precision-sensitivity points with minimum coverage threshold set to 0. The accession numbers of these samples are shown in Table 11. (b) The precision-sensitivity curves for multi-exon transcripts. (c,f) The average AUC, sensitivity and precision of total transcripts running with minimum coverage set to 0. The error bars show the s.d. (the same for other panels). Here (×10^4^) means the values shown in the figures are AUC values times 10^4^. (d,g) The average AUC, sensitivity and precision of multi-exon transcripts running with minimum coverage set to 0. (e,h) The average sensitivity and precision of single-exon transcripts running with minimum coverage set to 0. (i,k) The average AUC of total transcripts for strand-specific and non-strand-specific samples. (j,l) The average AUC of total transcripts for samples aligned to GRCh37 and GRCh38.

For StringTie2, AutoTuneX also outperforms the default settings in 46 out of 65 samples for the top-1 set. On average, AutoTuneX delivers a 2.3% higher AUC than the default settings (Fig. 1f). For the top-5 set, the best AUC of those five parameter vectors surpasses that of the default settings in 59 out of 65 samples, with an average AUC improvement of 8.4%. For the top-30 set, the best AUC of those thirty parameter vectors surpasses that of the default settings in all samples, with an average AUC improvement of 11.1%.

Fig. 1c,f show that for ENCODE65, AutoTuneX achieves higher AUC values by maintaining similar sensitivity while delivering higher precision compared to the default settings. For Scallop, the average precision is 15%, 18%, and 19% higher than the default settings for the top-1, top-5, and top-30 sets, respectively. For StringTie2, the average precision is 1.2%, 8.7%, and 13% higher. These results indicate that AutoTuneX enhances the performance of transcript assemblers, particularly in balancing sensitivity and precision tradeoffs. Compared to Scallop, AutoTuneX achieves less improvement with StringTie2, possibly due to StringTie2’s already well-tuned default settings or its smaller parameter search space.

AutoTuneX performs similar to the default settings in detecting multi-exon transcripts, with comparable average AUC values, sensitivity and precision (Fig. 1d,g). For single-exon transcripts, AutoTuneX obtains lower sensitivity but higher precision compared to the default settings (Fig. 1e,h). For Scallop, the average precision is 191%, 174%, and 198% higher for the top-1, top-5, and top-30 sets, respectively. For StringTie2, the average precision is 2.4%, 16%, and 31% higher. These results indicate that AutoTuneX tends to more aggressively filter out low-expression single-exon transcripts to enhance precision at the expense of sensitivity.

These 65 samples were collected from the ENCODE project that provide pre-computed read alignments [Shao and Kingsford, 2017]. The dataset includes 50 strand-specific samples and 15 non-strand-specific samples. Additionally, 30 samples are aligned to the human reference genome GRCh37, while 35 samples are aligned to GRCh38. Fig. 1i-l show that, overall, AutoTuneX outperforms the default settings for both Scallop and StringTie2, regardless of library type or reference genome.

The precision-sensitivity curves for the five samples showing the most significant AUC improvements with AutoTuneX are presented in Fig.1a, b for Scallop and in Fig.8a, b for StringTie2. For Scallop, each of these samples achieves more than an 80% increase in AUC using the parameter vectors recommended by AutoTuneX. For StringTie2, each sample achieves more than a 40% increase in AUC. In general, the precision-sensitivity curves generated by AutoTuneX consistently lie to the right of those from the default settings for both Scallop and StringTie2, whether considering total (single- and multi-exon) transcripts (Fig.1a and Fig.8a) or multi-exon transcripts only (Fig.1b and Fig.8b). These results further demonstrate the superior performance of AutoTuneX.

We also evaluate the performance of AutoTuneX on assemblers with different minimum coverage thresholds. The default minimum coverage thresholds for Scallop and StringTie2 are 1.01 and 1.0, respectively. The performance of AutoTuneX with the default minimum coverage thresholds (Fig.8c-h) is similar to its performance when the minimum coverage threshold is set to 0 (Fig.1c-h).

We compared AutoTuneX with the parameter advising method introduced by DeBlasio et al. [2020], which is the only previous method that explores parameter configuration for transcript assemblers. That approach generates a fixed set of 30 recommended parameter vectors to be used for all RNA-seq samples. To ensure a fair comparison, we compared that method with AutoTuneX’s top-30 set (Fig. 9). For Scallop, AutoTuneX outperforms the previous method in 56 out of 65 samples, with 18 samples achieving more than a 5% increase in AUC compared to the method in DeBlasio et al. [2020]. Similarly, for StringTie2, AutoTuneX outperforms the previous method in 56 out of 65 samples, with 22 samples achieving more than a 5% increase in AUC compared to the method in DeBlasio et al. [2020]. These results collectively highlight the strong performance of AutoTuneX.

We also compare AutoTuneX with two simpler methods on the ENCODE65 dataset:

- Direct application of the Bayesian optimization on each sample in ENCODE65.
- Implementation of a method called AutoMash, which follows the same framework as AutoTuneX but replaces the contrastive learning framework with the Mash distance [Ondov et al., 2016], a metric that measures the similarity between RNA-seq samples, to predict optimal parameters for new RNA-seq samples.

The results of these comparisons are displayed in Figs. 9-11. Generally, AutoTuneX has superior performance over both BO and AutoMash. These findings suggest that directly employing BO on each new sample is not the most efficient approach, and that the contrastive learning framework employed in AutoTuneX is more effective in predicting optimal parameters for transcript assemblers.

We further tested AutoTuneX on a larger RNA-seq dataset, referred to as SRA-test, from DeBlasio et al. [2020]. It contains 1588 RNA-Seq samples from the Sequence Read Archive. We again compared parameters predicted from AutoTuneX with the default parameter setting of the assembler and the method of DeBlasio et al. [2020] (Fig. 12). AutoTuneX outperforms the default settings for Scallop in terms of AUC in 81.9% of samples for the top-1 set, 97.0% for the top-5 set, and 99.0% for the top-30 set. On average, the AUC is improved by 12.0%, 21.5%, and 24.4% for the top-1, top-5, and top-30 sets, respectively (Fig.12e). AutoTuneX outperforms the approach of DeBlasio et al. [2020] in 95.3% of samples for the top-30 set, with an average AUC increase of 4.23% (Fig. 12e). These improvements are achieved by AutoTuneX through a slight reduction in sensitivity while enhancing precision (Fig. 12e).

For StringTie2, AutoTuneX outperforms the default setting in 84.5% of samples for the top-1 set, 97.7% for the top-5 set, and 98.8% for the top-30 set. On average, the AUC is improved by 2.36%, 4.83%, and 5.65% for the top-1, top-5, and top-30 sets, respectively (Fig.12f). AutoTuneX outperforms the method DeBlasio et al. [2020] in 97.7% of samples for the top-30 set, with an average AUC increase of 3.42% (Fig.12f). In this case, AutoTuneX achieves higher AUC values through a slight reduction in precision but higher sensitivity (Fig.12f).

We also present the precision-sensitivity curves for the five samples with the largest AUC improvements using AutoTuneX for both Scallop (Fig.12a,b) and StringTie2 (Fig.12c,d). The sample with the greatest AUC improvement for Scallop (SRR1177745) shows a 423% increase in AUC with the parameter vectors recommended by AutoTuneX, while the sample with the greatest improvement for StringTie2 (SRR1030497) shows a 674% increase. When considering total transcripts, all curves generated by AutoTuneX are clearly shifted to the right (Fig.12a,c). For multi-exon transcripts, many curves generated by AutoTuneX also similarly shift to the right (Fig.12b,d), demonstrating improved precision-sensitivity performance.

The most time-consuming part of training AutoTuneX for a specific transcript assembler is to search for the optimal parameter vectors for representative samples using Bayesian optimization. Training AutoTuneX for human RNA-seq samples requires approximately 15 days using around 300 CPUs to complete BO on all samples in ℛ_*rep*_. After AutoTuneX has been trained, it can be applied rapidly (*<* 5 minutes) to any new sample.

AutoTuneX provides a new strategy for automatically optimizing the use of sequence analysis, in particular transcript assemblers. Rather than developing new transcript assemblers, AutoTuneX improves the performance of existing methods by advising better parameters. Leveraging Bayesian optimization and contrastive learning, AutoTuneX uses a new BO-informed embedding framework for RNA-seq samples, positioning samples with similar optimal parameter vectors in close proximity. AutoTuneX demonstrates significant improvements in assembling human RNA-seq samples across different assemblers. The concept behind AutoTuneX can be naturally used for optimizing parameters of other bioinformatics tools.

## 2 Methods

### 2.1 Overview of AutoTuneX

Let ℛ be the universe of all possible RNA-seq samples and 𝒯 be the universe of all sets of transcripts. A transcript assembler, denoted as 𝒜, is a map 𝒜 (ℛ; *θ*) : ℛ × Θ → 𝒯 which takes a sample *R* ∈ ℛ and a parameter vector *θ* ∈ Θ ⊆ ℝ^*n*^ in some parameter space Θ as inputs and outputs a set of transcripts.

Given some loss function ℒ : 𝒯 → ℝ that estimates the performance of transcript assembly, we use *f* (*R*; *θ*) ≜ ℒ ∘ 𝒜 (*R*; *θ*) : ℛ × Θ → ℝ to estimate the performance of assembler 𝒜 with the input sample *R* and the parameter vector *θ*. We define *f*_*R*_(*θ*) ≜ ℒ (𝒜 (*R*; *θ*)) for cleaner notation. The objective in creating a parameter tuning system is to develop an oracle 𝒪 : ℛ → Θ, which can accurately determine an optimal or near-optimal parameter vector for each RNA-seq sample, that is:

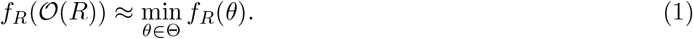

Our method, AutoTuneX, constructs such an oracle by first developing a mapping ℳ. This mapping takes an RNA-seq sample *R* as the input, and outputs an advisor set which is a set of good parameter candidates for this sample. Using this mapping, we then define the oracle 𝒪_ℳ_ as the following:

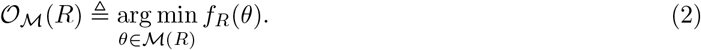

Different from traditional hyper-parameter tuning in machine learning, which focuses on optimizing hyper-parameters for a single specific function, ℳ operates as a meta-framework. This framework is able to minimize *f*_*R*_(*θ*) for each sample *R* by producing sample-specific advisor sets. A significant advantage of ℳ is that once it is trained, it does not require retraining or additional optimization for new samples, thus offering a more efficient and adaptable approach to parameter tuning across diverse samples.

As shown in Fig. 2 and described in the main text, training AutoTuneX is composed of three steps: (i) choose a representative sample set ℛ_*rep*_, from the sample universe ℛ; (ii) search for the best parameter vector for each representative sample 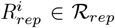 via Bayesian Optimization; and (iii) develop a similarity measure 𝕊 for RNA-seq samples through contrastive learning by using the information from step (ii). The culmination of these steps results in the definition of our mapping ℳ. Specifically, ℳ is defined as the union of the optimal parameter vectors from those representative samples that are identified as the nearest neighbors of a given sample *R* under the similarity measure 𝕊. That is:

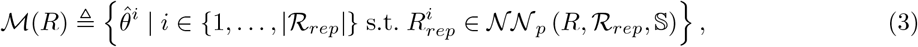

where 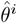 represents the optimal parameter vector for the representative sample 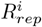 as determined through Bayesian Optimization in step (ii), and 𝒩𝒩_*p*_ (*R*, ℛ_*rep*_, 𝕊) is the set of *p* samples within ℛ_*rep*_ that are closest to the sample *R* under the measure 𝕊. We now use the following three sections to describe the three steps (i) to (iii) in detail.

**Figure 2:**
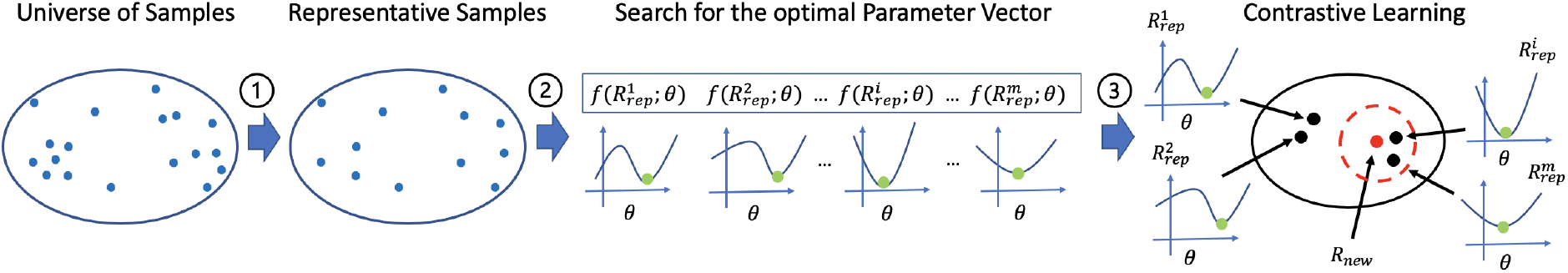
Overview of AutoTuneX. Step 1: Representative sample selection from ℛ, each blue dot represents one human RNA-seq sample, and the blue ellipse represents the universe of all human RNA-seq samples; Step 2: Search for the best parameters *θ* for representative samples via Bayesian optimization, *f* represents a function quantifying the performance of parameter settings for a specific sample, and each green dot represents the optimal parameter point; Step 3: Learning a embedding framework via contrastive learning. Samples with similar optimal parameters are mapped close. The dashed red circle represents the region of the nearest neighbors of a new sample *R*_*new*_ in the latent space.

### 2.2 Representative sample selection from ℛ

To construct ℳ, it is essential to use existing RNA-seq samples as the training foundation. The training set needs to effectively encapsulate the critical transcriptional characteristics of ℛ, the entire spectrum of RNA-seq samples, to ensure that ℳ possesses strong generalizability to new samples. One approach could be to incorporate all existing RNA-seq samples into the training set. However, considering the immense volume of available RNA-seq data – numbering in the hundreds of thousands of samples – such an approach would demand prohibitively high computational resources. To strike a balance between comprehensive representation and computational feasibility, we adopt the representative sample set identified in Tung and Kingsford [2021], which includes around 7,000 human RNA-seq samples. This set effectively represents the broader collection of existing human RNA-seq samples. To enhance the computational efficiency, we further use apricot [Schreiber et al., 2020], for selecting a smaller representative subset from these samples. This selection process ensures a robust coverage of ℛ, while significantly alleviating computational demands. The final subset of representative samples selected through this method is denoted as ℛ_*rep*_.

### 2.3 Search for the best parameter vectors for representative samples via warm-up-based Bayesian Optimization

For every representative sample 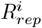 in ℛ_*rep*_, we use BO to search for its best parameter vector for a particular assembler by minimizing the function 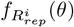. BO is particularly suitable for this task for two main reasons: (i) 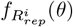 is essentially a black-box function, meaning its derivatives are difficult to compute. This complexity arises because the transcript assembler 𝒜 typically comprises multiple modules that tackle intricate combinatorial problems. As a result, it is not feasible to express 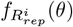 in a closed form; and (ii) each evaluation of 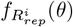 necessitates a run of the transcript assembler 𝒜 which can be a slow process, ranging from minutes to hours. Therefore, even a single evaluation of 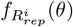 can be quite time-consuming.

However, standard Bayesian Optimization algorithms are not immediately applicable in our context due to their reliance on a predefined search domain. Typically, these algorithms require a user-specified domain, assumed to encompass the optimal or near-optimal points for the function being optimized [Shahriari et al., 2016]. In the case of RNA-seq samples, however, the challenge arises from the variability of optimal parameter vectors across different samples. Given this diversity, and the potential for significant dissimilarities between optimal parameters for various samples, it becomes impractical to establish a fixed search domain suitable for all cases. This aspect of the RNA-seq samples necessitates an adaptation or modification of standard Bayesian Optimization techniques to accommodate the varying nature of the optimal parameter vectors.

Inspired by Poloczek et al. [2016] and DeBlasio et al. [2020], we propose a new warm-up-based BO, called CAWarm-BO (<u>C</u>oordinate <u>A</u>scent <u>Warm</u>-up <u>B</u>ayesian <u>O</u>ptimization), that is able to automatically adjust the search domain for each sample. The effectiveness of this new framework is based on the following observations:

- Coordinate ascent is an iterative optimization algorithm that successively maximizes along the coordinate directions of a function to find the maximum [Zangwill, 1969]. Coordinate ascent does not require a predefined search domain, and it will rapidly converge towards a local region potentially containing the optimal or near-optimal points. However, within that localized region, coordinate ascent tends to be slower in pinpointing the local optimal point.
- Bayesian Optimization is able to efficiently find the best point within a reasonably sized search domain, but the search domain must be predefined.

By synergizing these two methodologies, CAWarm-BO leverages the strengths of both: using coordinate ascent to swiftly approach a promising region without the need for a predefined domain, followed by Bayesian Optimization to efficiently find the optimal point within that localized area.

We run CAWarm-BO independently on each sample in the representative set ℛ_*rep*_ (here, we maximize 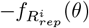 which is equivalent to minimizing 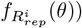. For each function 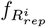, CAWarm-BO yields a set of pairs denoted as 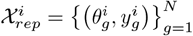. In this set, 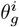 represents a queried parameter vector during the Bayesian Optimization search, and 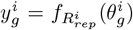 corresponds to the loss value for that parameter vector. The optimal parameter vector 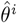 for sample 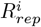 is determined as the parameter vector 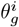 in 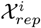 that yields the smallest 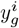.

### 2.4 Learning the similarity measure via contrastive learning

Compared to other optimization algorithms, CAWarm-BO is more efficient in optimizing the function *f*_*R*_(*θ*) for a given sample *R*. However, as shown in our experimental results, it is still not efficient enough that we can directly use it to search for the best parameter vector for every new sample, i.e. it cannot be solely relied upon as the oracle 𝒪 due to its slow speed. This limitation underscores the need to develop a similarity measure 𝕊, enabling model ℳ to quickly generate the specific advisor set for each new sample through nearest neighbor search, as shown in Eq. 3. We learn 𝕊 via the contrastive learning (CL) framework. Within this framework, given any two RNA-seq samples *R*_1_ and *R*_2_, they are embedded into a latent space via a feature encoder *Enc*_*ϕ*_ with trainable parameters *ϕ*. In this latent space, the similarity between two vectors is quantified using the cosine similarity measure:

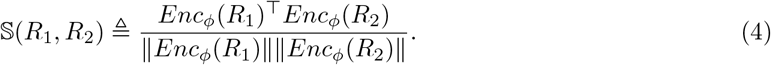

Learning 𝕊 is now equivalent to training the encoder *Enc*_*ϕ*_. Our training framework is inspired by previous work of supervised contrastive learning such as Khosla et al. [2020] and Wang et al. [2022], but has the following several unique components:

- We represent each sample in ℛ_*rep*_ by a set derived from the MinHash sketch [Broder, 1997, Ondov et al., 2016]. These sets serve as the inputs for the encoder *Enc*_*ϕ*_. The details are described in Section 2.4.1.
- We develop a novel method to define the similarities between pairs of representative samples, denoted as 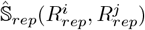 for training purposes. The details are described in Section 2.4.2.
- We develop a distinctive data augmentation module for enhancing the robustness and accuracy of training. The details are described in Section 2.4.3.
- We develop a neural network architecture and a contrastive loss function that are particularly tailored to our specific requirements. The details are described in Section 2.4.4.

The pseudocode of the training framework is given in Algorithm 1. During each training iteration, every input sample in the training batch is embedded using the encoding network *Enc*_*ϕ*_ (line 5 of Alg. 1). The outputs from *Enc*_*ϕ*_, together with 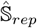, are used to compute the contrastive loss function (ℒ_*CL*_) (line 7 of Alg. 1). We train *Enc*_*ϕ*_ by minimizing ℒ_*CL*_ (line 8 of Alg. 1) via gradient-based algorithms.

#### Algorithm 1 Contrastive Training Module

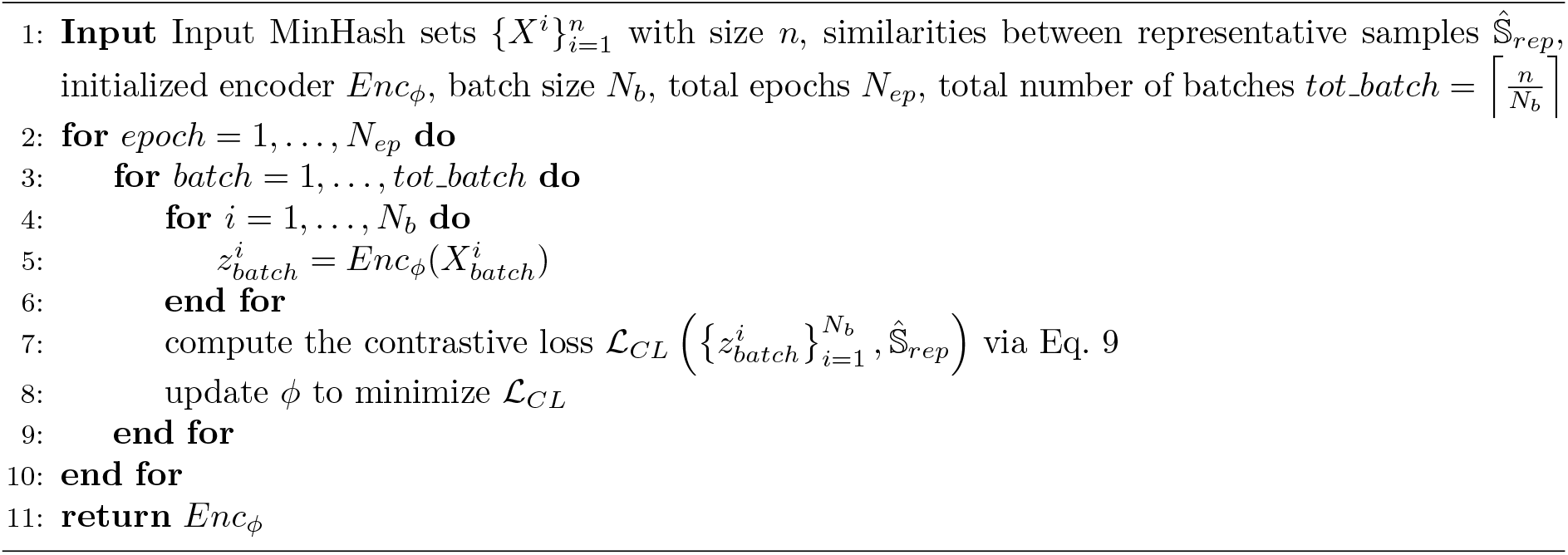

To obtain an advisor set for a new sample, we use the learned *Enc*_*ϕ*_ to map this new sample to the embedding space (line 3 of Alg. 2) and use the cosine similarity to identify the closest neighbors among the representative samples (lines 6 of Alg. 2). The advisor set is then composed of the best parameter vectors from these nearest neighbors (lines 8 of Alg. 2).

#### Algorithm 2 Nearest Neighbor Computation for a New Sample

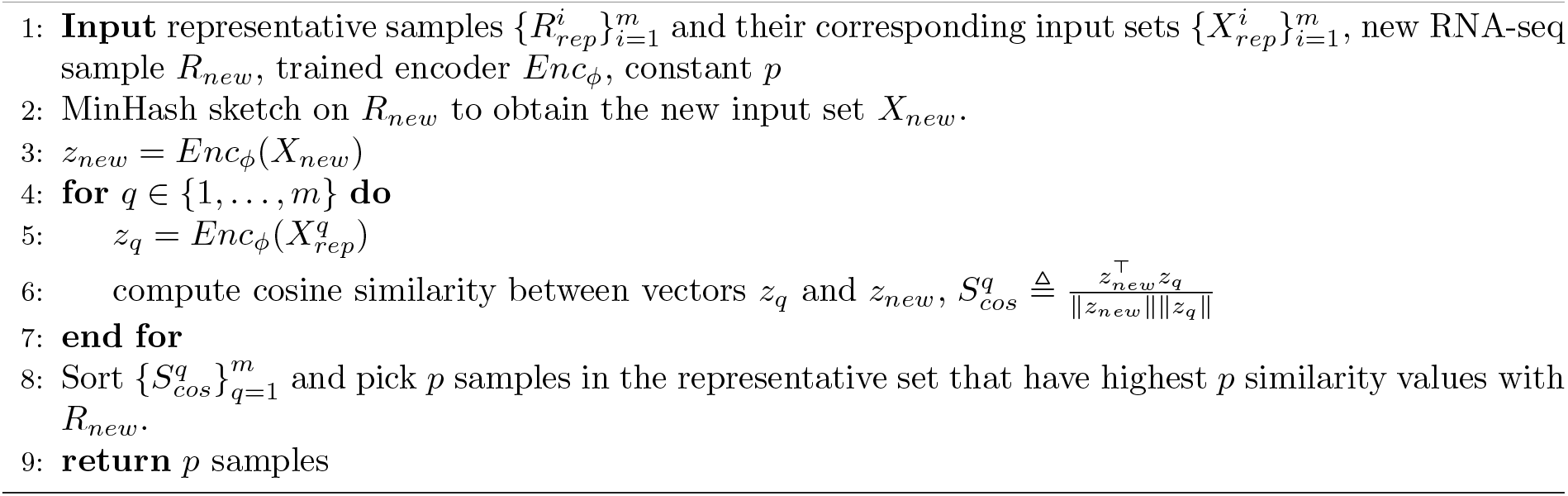

#### 2.4.1 RNA-seq sample representation

Given that each RNA-seq sample comprises a vast number of sequencing reads, which are not ideally suited for direct input into neural networks, we use the MinHash sketch [Broder, 1997, Ondov et al., 2016] to generate representations for these samples. MinHash sketch process begins by decomposing the reads in an RNA-seq sample into constituent k-mers. Each k-mer is then processed through a hash function *h* to obtain a hash value. Finally, for a predetermined sketch size *s*, MinHash sketch returns the *s* smallest hashes output by *h* over all k-mers, along with the count of k-mers associated with these hash values. Each RNA-seq sample *R* is represented by a set input 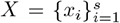, where each *x*_*i*_ is a two-dimensional vector; the first dimension denotes a hash value, and the second dimension indicates the number of k-mers in *R* corresponding to this hash.

#### 2.4.2 Similarity between representative samples

Contrary to supervised contrastive learning in classification problems [Khosla et al., 2020] where sample pairs are simply categorized as positive (having the same label) or negative (having different labels), our case more closely aligns with a contrastive regression problem as in Wang et al. [2022] where 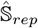 is used to assess the degree of similarity between two RNA-seq samples. Ideally, for each pair of representative samples) 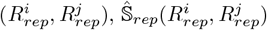 should correspond to the similarity between their respective functions 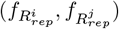 However, since these functions lack a closed-form representation, direct computation of their similarity is not feasible. Instead, we leverage information from the Bayesian Optimization (BO) step to approximate these functions and assess their similarity. Specifically, as described above, the output of CAWarm-BO for each sample 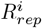 is a set of pairs 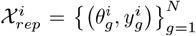. We use Gaussian Process (𝒢 𝒫) models to fit data: 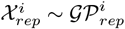 and 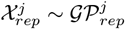 The posterior means of the Gaussian Process models, denoted as 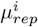 and 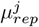 respectively, are estimations of functions 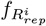 and 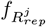 We then introduce the following term, called *normrank*, to quantify the similarity between 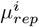 and 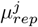

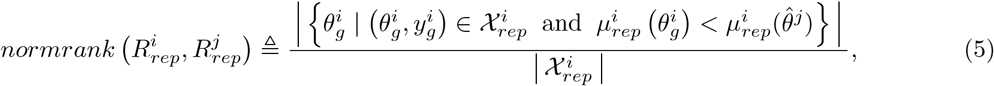

where 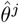 is the optimal parameter vector for 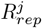 found by BO. The numerator of Eq. (5) represents the number of parameter vectors queried in 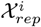 that have smaller posterior mean values than the value of the optimal parameter vector for 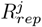 Intuitively, if 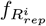 and 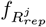 are similar, the optimal parameter vector for 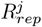 should also be an approximately optimal one of 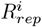, making the numerator smaller. The denominator of Eq. 5 serves as a normalization term, accounting for different sizes of 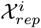 *normrank* functions as a distance-like measure between samples: the more alike the samples are, the lower the *normrank* value. However, it has two drawbacks: (i) it is not symmetric, meaning *normrank* 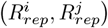, may not be equal to *normrank*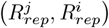, and (ii) it does not constitute a true distance as it fails to satisfy the triangle inequality. To address these issues, we first compute the average 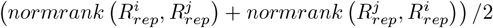 to ensure symmetry. We then apply the shortest-path method, as introduced in Lesne et al. [2014], to reconstruct the distances, denoted as 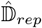, and to filter out some noise. Based on this, 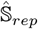 is subsequently defined as:

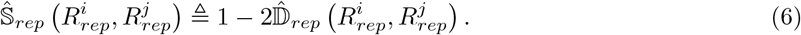

According to Eq. (5), for any pair of samples, the value of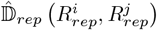 falls within the range [0, 1]. Consequently, as per Def. (6), this ensures that the value of 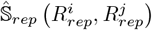 lies within the range [−1, 1], aligning with the range typical for cosine similarity.

#### 2.4.3 Data augmentation module

The time-intensive nature of conducting Bayesian Optimization (BO) on each RNA-seq sample to acquire the optimal parameter vector imposes a constraint on the size of the representative sample set; it cannot be excessively large. This limits the amount of training data available for contrastive learning. As will be discussed in Section 3.2, the representative sample set comprises around 1,000 samples. To improve the accuracy of training under these constraints, we develop a novel data augmentation method. This method substantially expands the size of the training dataset without necessitating additional rounds of BO computation.

The underlying principle of our data augmentation method is as follows: a new RNA-seq sample *R*_*sub*_ is created by subsampling reads from an original sample *R*. Its corresponding function 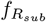 is expected and *R* to have a similar optimal parameter vector to that of *f*_*R*_. This means we can create additional training data by subsampling reads from the original RNA-seq samples, and we can calculate the similarity scores without the need for extra Bayesian Optimization (BO) computations. Specifically, let 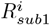 and 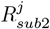 be subsampled versions of 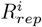 and 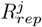 respectively. The similarity score between these two new subsampled samples is defined as follows:

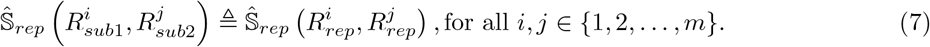

In practice, for each representative sample, we generate seven subsampled variants with sampling ratios 0.65, 0.7, 0.75, 0.8, 0.85, 0.9, 0.95. This approach effectively increases the size of our training data by a factor of seven, without necessitating any additional Bayesian Optimization (BO) computations. The effectiveness of this data augmentation module is further discussed in Section 3.3.3.

#### 2.4.4 Neural network architecture and loss function

As outlined in Section 2.4.1, each RNA-seq sample is represented as a set. Consequently, in our case, it is necessary to use a neural network architecture that is capable of processing sets as inputs. Crucially, this architecture must exhibit permutation invariance, meaning its output should remain unchanged irrespective of the order of elements within the set.

Inspired by Zaheer et al. [2017], the encoder *Enc*_*ϕ*_ is designed to be a composite of two sub-networks: *h*_*ϕ*_ and *g*_*ψ*_ with trainable parameters *ϕ* and *ψ*. Specifically, given an RNA-seq sample *R* with its set representation 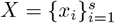, we have:

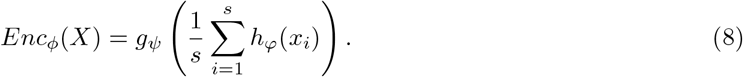

Theorem 2 of Zaheer et al. [2017] proved that the form of Eq. (8) preserves permutation invariance. Our contrastive loss function is simply defined as mean squared error:

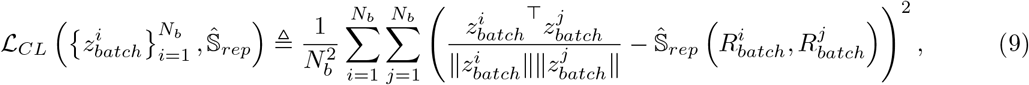

where *N*_*b*_ is the batch size and 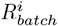 and 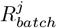 are RNA-seq samples in the training data corresponding to embeddings 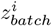 and 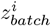. Compared to the contrastive loss formulas used in Khosla et al. [2020] and Wang et al. [2022], we find that this simple loss function achieves very good performance in our case, with the advantage that no extra hyper-parameters such as the temperature need to be tuned.

##### Code availability

AutoTuneX is implemented in Python and is freely available as open source software at https://github.com/Kingsford-Group/autotunex.

## 3 Supplementary Material

### 3.1 Additional Related Work

#### 3.1.1 Parameter tuning in bioinformatics

Parameter tuning has been previously discussed in several bioinformatic areas such as multiple sequence alignment [Cedillo et al., 2022, DeBlasio and Kececioglu, 2014, 2015b,a], biological pathway reconstruction [Magnano and Gitter, 2021] as well as transcript assembly [DeBlasio et al., 2020]. The method proposed in DeBlasio et al. [2020] is the first and only work developing a parameter tuning system for transcript assembly by learning the advisor set in a data-driven way. Although the method achieves significantly better performance than the default settings of transcript assemblers on Scallop [Shao and Kingsford, 2017] and StringTie [Pertea et al., 2015], the method is limited by its reliance on a fixed advisor set for all new samples. This limitation is compounded by the small number of RNA-seq samples used in its training set, which fails to comprehensively represent the diversity of all RNA-seq samples. Consequently, this approach does not work well when encountering new samples markedly different from those in the training set, a common issue with small training datasets.

AutoTuneX addresses these shortcomings by employing a large, representative collection of RNA-seq samples. This allows for the learning of a system capable of generating sample-specific advisor sets, thereby offering a more adaptable and robust solution in the context of transcript assembly.

#### 3.12 Bayesian optimization

Bayesian optimization (BO) is a sample-efficient framework designed for globally optimizing a black-box function *f* (*x*) that is costly to evaluate [Frazier, 2018, Shahriari et al., 2015]. It aims to obtain the input point that maximizes (or minimizes) the function *f* by iteratively acquiring input points that are likely to achieve the maximum and evaluating the function on these points. Each iteration of BO has two components: a Bayesian statistical model, usually a Gaussian Process regressor [Schulz et al., 2018], to estimate *f* (*x*), and an acquisition function that determines the next input point to sample. We refer the reader to Frazier [2018] for a detailed description of BO. Although BO has been successfully applied in many scenarios such as hyper-parameter tuning [Snoek et al., 2012, Klein et al., 2017], reinforcement learning [Brochu et al., 2010, Marco et al., 2017, Wilson et al., 2014], and chemical design [Griffiths and Hernández-Lobato, 2020, Negoescu et al., 2011], its efficiency in tuning parameters of bioinformatics tools has not been broadly explored.

In our study, we introduce the use of Bayesian Optimization (BO) as an effective method for determining the optimal transcript assembly parameters for a specific RNA-seq sample. We use a transcript assembler paired with a loss function designed to estimate the assembler’s performance. In this scenario, the task of finding the best parameters for a given RNA-seq sample *R* is equivalent to optimizing the loss function. As we demonstrate below, this loss function manifests as a black-box function, hence BO emerges as a highly suitable tool for this optimization process due to its proficiency in handling such black-box scenarios.

#### 3.1.3 Contrastive learning

Contrastive Learning (CL) is a class of learning methods that learns by comparing input samples. The goal of CL is to learn embeddings of input samples such that those of “similar” samples (e.g. two cat images) are embedded close together, while those of “dissimilar” samples (e.g. one cat image and one dog image) are placed further away [Le-Khac et al., 2020]. A typical CL framework has three components:

- Similarity definition: this determines whether a pair of samples in the training set is similar (positive pair) or dissimilar (negative pair). The criteria for similarity vary depending on the specific problem. For instance, in supervised learning, the labels of input samples might be used to define similarity: samples sharing a label are considered similar, and those with different labels are dissimilar [Khosla et al., 2020].
- Feature encoder: a trainable neural network that maps each input sample into an embedding space.
- Contrastive loss function: this function, using the outputs of the feature encoder and the defined similarity, guides the training process. By minimizing this loss, the encoder learns to map samples from a positive pair close together and those from a negative pair far apart in the embedding space.

CL has been effectively applied in various domains such as self-supervised learning [Chen et al., 2020, Chuang et al., 2020], image classification [Khosla et al., 2020], gaze direction regression [Wang et al., 2022] and metagenomics [Zhang et al., 2022b]. In this work, we adapt CL to compare different RNA-seq samples. We define “similarity” in the context of RNA-seq samples (details below) based on the premise that similar samples will likely have similar optimal parameter vectors for transcript assembly. Thus, knowing the optimal parameters for one RNA-seq sample allows us to apply them to a similar sample, bypassing the need for fresh optimization.

### 3.2 Experimental Details

We test AutoTuneX on Scallop v0.10.2 (𝒜 = Scallop) with 18 tunable parameters and StringTie2 v2.2.1 (𝒜 = StringTie) with 8 tunable parameters. The names of these parameters, their default settings and their types are listed in Table 1 and Table 2.

**Table 1:** Parameter information of Scallop.

| Parameter | Default value | Type |
| --- | --- | --- |
| uniquely_mapped_only | False | binary |
| use_second_alignment | False | binary |
| max_dp_table_size | 10000 | integer |
| max_edit_distance | 10 | integer |
| max_num_exons | 1000 | integer |
| min_bundle_gap | 50 | integer |
| min_exon_length | 20 | integer |
| min_flank_length | 3 | integer |
| min_mapping_quality | 1 | integer |
| min_num_hits_in_bundle | 20 | integer |
| min_router_count | 1 | integer |
| min_splice_boundary_hits | 1 | integer |
| min_subregion_gap | 3 | integer |
| min_subregion_length | 15 | integer |
| min_transcript_length_base | 150 | integer |
| min_transcript_length_increase | 50 | integer |
| max_intron_contamination_coverage | 2 | float |
| min_subregion_overlap | 1.5 | float |

**Table 2:** Parameter information of StringTie2.

| Parameter | Default value | Type |
| --- | --- | --- |
| t | False | binary |
| u | False | binary |
| m | 200 | integer |
| a | 10 | integer |
| g | 50 | integer |
| j | 1 | float |
| f | 0.01 | float |
| M | 1 | float |

During AutoTuneX’s training, we use the area under the curve (AUC) of an assembler’s prediction of known transcripts to evaluate the performance of the two transcript assemblers. AUC is a widely used metric for assessing transcript assembler performance [Shao and Kingsford, 2017, DeBlasio et al., 2020], as it provides a single value that balances sensitivity and precision tradeoffs. In this context, we first use the standard tool gffcompare [Pertea and Pertea, 2020] to compute the accuracy of the predicted transcripts: a predicted multi-exon transcript is considered as correct if its intron chain can be exactly matched to a (multi-exon) transcript in the transcript assnotation file, while a predicted single-exon transcript is defined as correct if it overlaps at least 80% with a (single-exon) transcript in the annotation file. Gffcompare then reports two metrics: sensitivity (the ratio between the number of correct transcripts and the total number of transcripts in the annotation file) and precision (the ratio between the number of correct transcripts and the total number of predicted transcripts). By ranking the predicted transcripts according to their expression abundance, we draw the precision-sensitivity curve and calculate the AUC value as the area under this curve. The loss function L for training is defined as the negative AUC value.

During testing, we also evaluate the performance of AutoTuneX using two additional metrics: the total number of correct transcripts (which is proportional to sensitivity) and precision, in addition to AUC. Although these metrics may underestimate accuracy—since novel transcripts that are correctly assembled are considered incorrect due to their absence in the current annotation file—they have been widely used in previous studies [Pertea et al., 2015, Song et al., 2016, Liu et al., 2016, Shao and Kingsford, 2017, Kovaka et al., 2019] for comparing transcript assemblers, as they effectively reflect the relative accuracy of different assemblers.

The transcript annotation file used for training is GENCODE release 24 [Frankish et al., 2019], while GENCODE release 46 is used for testing to ensure that the model does not overfit to a specific annotation version.

Since transcript assemblers including Scallop and StringTie2 usually have mixed types of parameters (parameters include categorical, integer, and continuous variables), we choose CASMOPOLITAN [Wan et al., 2021], a BO-based algorithm for mixed search spaces, as the basic framework of CAWarm-BO. The initial search domain for each non-binary parameter is defined as [0, 10 × default] (e.g. for max dp table size in Table 1, the initial range includes all integers within [0, 100000]). CAWarm-BO initially runs the same coordinate ascent algorithm as proposed by DeBlasio et al. [2020], with a maximum warmup iteration set at 60. The step size for each coordinate in the ascent is determined by the initial step size specified by DeBlasio et al. [2020]. The warmup phase of CAWarm-BO will cease if it exhaustively interrogates the entire parameter set without any improvement or upon reaching the maximum warmup iteration.

Assuming the best parameter vector identified during warmup is 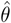, where 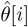 represents the value of its *i*-th dimension, CAWarm-BO adjusts the search range of the *i*-th dimension to [0, max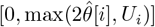], with [0, *U*_*i*_] being the initial range of the *i*-th dimension. Following this, CAWarm-BO executes CASMOPOLITAN over this new domain. The total number of iterations for CAWarm-BO per sample is set to 200, inclusive of the warmup phase.

From human samples in Sequence Read Archive [Cochrane et al., 2016], 7, 860 human RNA-seq samples are selected by Tung and Kingsford [2021] as representative samples. We then use apricot [Schreiber et al., 2020] to further select 1, 500 samples for constructing ℛ_*rep*_. These 1, 500 samples are aligned using the aligner STAR [Dobin et al., 2013] to the human reference genome GRCh38. We run CAWarm-BO independently on these samples, and remove samples that have very low AUC values (the samples for which the best AUC value found by BO is less than 3 ×10^−4^). This filtration process resulted in 1,263 samples retained for contrastive training for Scallop, and 1,235 samples for StringTie2.

As described in Section 2.4.1, we use MinHash sketch to create the set input 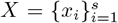 for each RNA-seq sample *R*. Each value in each *x*_*i*_ is z-score normalized. We set *k* = 21 and *s* = 1000 which are default settings in Ondov et al. [2016] and exclude *k*-mers with counts less than 2.

We use PyTorch [Paszke et al., 2019] to implement the contrastive learning framework. The architecture of *h*_*ϕ*_ is a fully connected neural network with layer sizes (2, 128, 128, 128, 128). Meanwhile, the architecture of the network *g*_*ψ*_ is another fully connected neural network with layer sizes (128, 256, 256, 256, 64). For each layer except the last one in both networks, we use the ReLU activation function, coupled with batch normalization [Ioffe and Szegedy, 2015] preceding each activation. We use ADAM [Kingma and Ba, 2014] with a learning rate 1 ×10^−3^ to optimize the neural network. The batch size in each iteration is 128 and the total number of epochs is 400. The training is conducted using one NVIDIA GeForce RTX 3080 Ti GPU.

As in DeBlasio et al. [2020], we use the following three datasets in our experiments:

- ENCODE10, a collection of 10 RNA-seq samples from the ENCODE database [The ENCODE Project Consortium, 2012]. These samples are already aligned in DeBlasio et al. [2020] to the human reference genome GRCh38. DeBlasio et al. [2020] use this dataset to generate the advisor set, while we use it to test the performance of CAWarm-BO.
- ENCODE65, a collection of 65 RNA-seq samples also from ENCODE. None of them is included in the representative sample set (training data). These samples have pre-existing alignments in ENCODE. We use them to test the performance of AutoTuneX.
- SRA-test, a collection of 1595 RNA-Seq samples from DeBlasio et al. [2020]. Among these samples, we find that two of them cannot be downloaded from the Sequence Read Archive [Cochrane et al., 2016] (ERR313180 and ERR313182). The remaining samples are aligned using the aligner STAR [Dobin et al., 2013] to the human reference genome GRCh38. We remove five samples are also in the representative sample set (SRR1030499, SRR1286920, SRR1812365, SRR496593, SRR592581), and then use the remaining 1588 samples to test the performance of AutoTuneX.

### 3.3 Additional experimental results

#### 3.3.1 Performance of CAWarm-BO

We compare CAWarm-BO with CASMOPOLITAN (without warmup and search domain adjustment) and coordinate ascent (CA) proposed in DeBlasio et al. [2020] on the ENCODE10 dataset. Fig. 3(a) and Fig. 4(a,b,c) show that in most samples, CAWarm-BO’s performance closely mirrors CA during the warmup phase but surpasses CA following the commencement of BO. CASMOPOLITAN shows competitive results in many samples when compared with CAWarm-BO, but without a warmup phase, it initially spends several iterations identifying a suitable local search region, resulting in slower convergence within the first 50 iterations. Furthermore, lacking search domain adjustment, CASMOPOLITAN sometimes only achieves a sub-optimal point, as seen in the middle panel of Fig. 3. We also evaluated the highest AUC values obtained by CAWarm-BO and CASMOPOLITAN within 200 iterations against the best values achieved by CA without any iteration limit (continuing CA until the stopping criteria set by DeBlasio et al. [2020] are met). Table 3 reveals that the best AUC values discovered by CAWarm-BO and CA are quite similar, although CA requires significantly more iterations than CAWarm-BO to reach these optimal values. Additionally, Table 4 indicates that this trend is consistent across different transcript assemblers. These findings indicate that CAWarm-BO is a more efficient method than both CA and CASMOPOLITAN for finding the best parameter vector for each sample in the representative sample set.

**Table 3:** Comparisons of Coordinate Ascent (CA), CASMOPOLITAN, and CAWarm-BO on ENCODE10 dataset. The transcript assembler used here is Scallop. Both CASMOPOLITAN, and CAWarm-BO are run within 200 iterations, while we run Coordinate Ascent until the stopping requirement set by DeBlasio et al. [2020] is satisfied. The table shows the best AUC value (×10^4^) found for each method and the number of iterations (in bracket) used for finding the best value.

| Samples | CAWarm-BO | CASMOPOLITAN | CA |
| --- | --- | --- | --- |
| SRR307903 | 611.70 (165) | 610.98 (146) | 613.12 (658) |
| SRR307911 | 545.20 (156) | 543.35 (198) | 549.39 (736) |
| SRR315323 | 411.25 (176) | 409.61 (150) | 409.29 (458) |
| SRR315334 | 576.45 (173) | 576.37 (156) | 579.69 (460) |
| SRR387661 | 298.49 (177) | 299.53 (188) | 296.57 (517) |
| SRR534291 | 534.00 (171) | 513.68 (200) | 534.17 (458) |
| SRR534307 | 732.26 (196) | 729.79 (182) | 733.01 (639) |
| SRR534319 | 318.37 (185) | 317.87 (96) | 315.98 (290) |
| SRR545695 | 395.08 (180) | 388.43 (80) | 393.33 (447) |
| SRR545723 | 551.56 (174) | 552.10 (144) | 552.81 (372) |

**Table 4:** Comparisons of Coordinate Ascent (CA), CASMOPOLITAN, and CAWarm-BO on ENCODE10 dataset. The transcript assembler used here is StringTie2. Both CASMOPOLITAN, and CAWarm-BO are run within 200 iterations, while we run Coordinate Ascent until the stopping requirement set by DeBlasio et al. [2020] is satisfied. The table shows the best AUC value (×10^4^) found for each method and the number of iterations (in bracket) used for finding the best value.

| Samples | CAWarm-BO | CASMOPOLITAN | CA |
| --- | --- | --- | --- |
| SRR307903 | 542.26 (186) | 541.50 (115) | 542.59 (387) |
| SRR307911 | 478.17 (191) | 478.52 (173) | 474.37 (328) |
| SRR315323 | 361.80 (174) | 358.15 (149) | 361.74 (289) |
| SRR315334 | 488.93 (195) | 489.20 (163) | 489.54 (302) |
| SRR387661 | 441.58 (182) | 441.57 (165) | 442.02 (274) |
| SRR534291 | 465.90 (165) | 463.69 (140) | 466.04 (225) |
| SRR534307 | 634.86 (173) | 635.65 (166) | 635.94 (316) |
| SRR534319 | 259.91 (197) | 260.39 (174) | 262.69 (304) |
| SRR545695 | 338.88 (181) | 333.90 (168) | 338.85 (303) |
| SRR545723 | 501.54 (178) | 500.47 (162) | 502.03 (292) |

**Figure 3:**
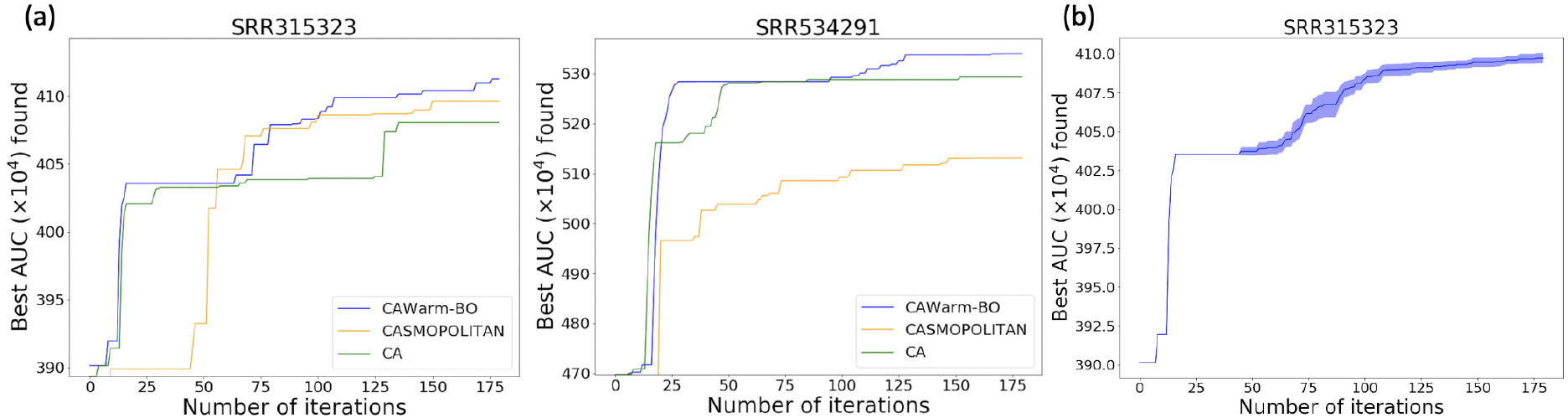
(a) Comparisons of Coordinate Ascent (CA), CASMOPOLITAN and CAWarm-BO on two RNA-seq samples from ENCODE10 (accession numbers: SRR315323 and SRR534291). The transcript assembler used here is Scallop. For each sample, we do one run for each method and plot the maximum value found by iterations. (b) The variance of CAWarm-BO with different runs. Here, we do 10 independent runs on the sample SRR315323, and plot the mean and 1*/*2 standard deviation of the maximum value found by iterations. Note that the warm-up part is deterministic.

**Figure 4:**
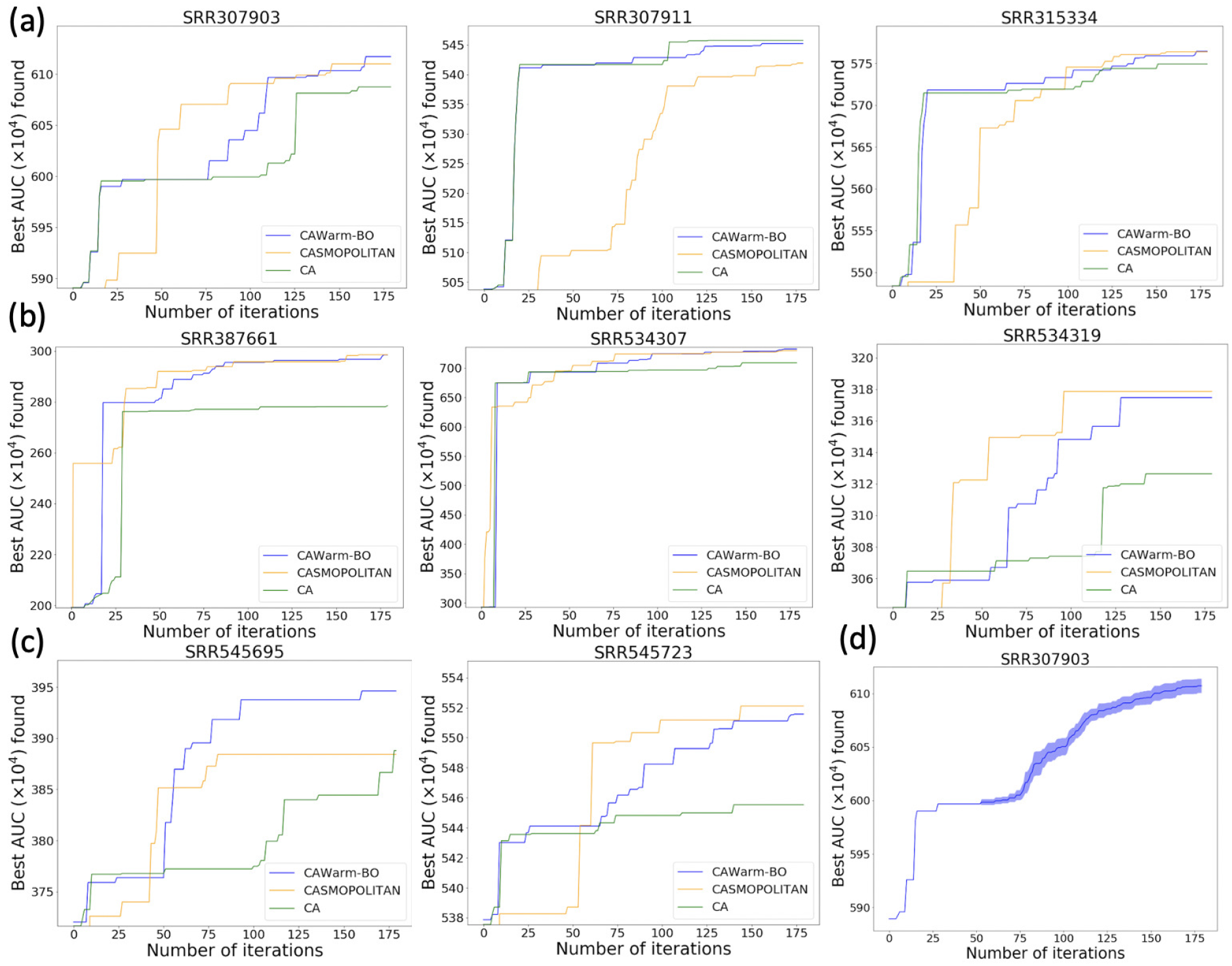
(a,b,c) Comparisons of Coordinate Ascent (CA), CASMOPOLITAN, and CAWarm-BO on eight RNA-seq samples from ENCODE10. The transcript assembler used here is Scallop. For each sample, we do one run for each method and plot the maximum value found by iterations. (d) The variance of CAWarm-BO with different runs. Here, we do 10 independent runs on the sample SRR307903, and plot the mean and 1*/*2 standard deviation of the maximum value found by iterations. Note that the warm-up part is deterministic.

#### 3.3.2 Accuracy of similarity definition between representative samples

We use the following correlation test to verify the accuracy of similarity values 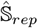 defined in Section 2.4.2 between representative samples. We randomly select 10 samples in the representative set. For each of these samples, denoted *R*, we compute *f*_*R*_(*θ*) using the best parameter vectors 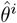 found by CAWarm-BO for all other samples in the representative set. We plot the relationships between these function values 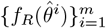 and the similarity values 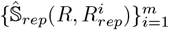, shown in Fig. 5(c). We compare 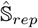 with two alternative similarity definitions: (i) Mash distance [Ondov et al., 2016], a distance metric based on the set representations of two samples. We calculate this distance between sample *R* and all others, plotting the function values against the Mash similarity (negative Mash distance values), shown in Fig. 5(a); (ii) The Euclidean distance between the best parameter vector of *R* (found by CAWarm-BO) and those of all other samples, normalized to the [0,1] range to negate scale effects. The relationship between the function values and the Euclidean similarity (negative Euclidean distance values) is also plotted, shown in Fig. 5(b). An ideal similarity measure should exhibit a strong negative correlation with these function values. Table 5 and Table 6, along with Fig. 5 show that in most tested samples 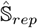 demonstrates a higher negative correlation with the function values compared to both Mash and Euclidean similarities, irrespective of the transcript assembler used. These results suggest that leveraging information from the BO steps enables more accurate quantification of sample similarity than directly computing the distance between set representations of RNA-seq samples. Moreover, the *normrank* function used in 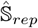 incorporates broader information from the BO step than just the best parameter vector of each sample, thus enhancing the accuracy of similarity quantification.

**Table 5:** Spearman’s rank correlation coefficient between similarity measures and function values for different samples. The transcript assembler used here is Scallop.

| Samples | Mash similarity | Euclidean similarity | $\hat{S}_{rep}$ |
| --- | --- | --- | --- |
| ERR197922 | -0.153 | -0.112 | **-0.502** |
| SRR1812365 | -0.460 | -0.277 | **-0.600** |
| SRR5099917 | -0.353 | 0.114 | **-0.664** |
| SRR492054 | -0.713 | -0.363 | **-0.891** |
| SRR346466 | -0.619 | -0.682 | **-0.842** |
| SRR6687435 | -0.372 | -0.512 | **-0.754** |
| SRR1782845 | -0.556 | -0.480 | **-0.757** |
| SRR8535251 | -0.327 | -0.238 | **-0.435** |
| SRR3932352 | -0.627 | -0.665 | **-0.894** |
| ERR2619102 | -0.254 | 0.348 | **-0.432** |

**Table 6:**
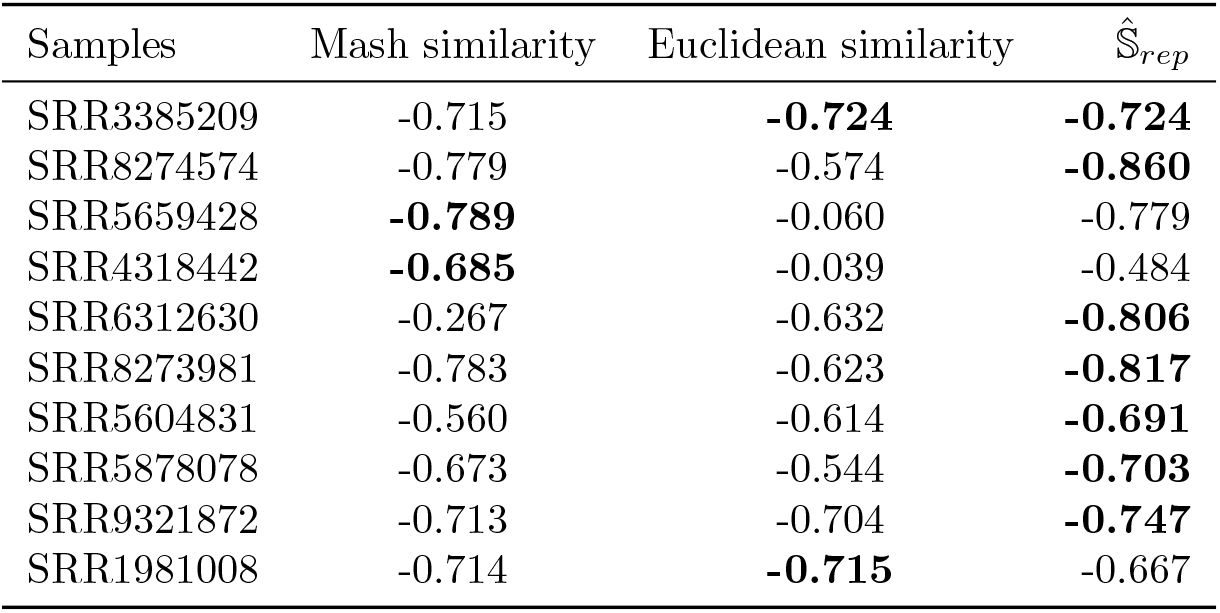
Spearman’s rank correlation coefficient between similarity measures and function values for different samples. The transcript assembler used here is StringTie2.

**Figure 5:**
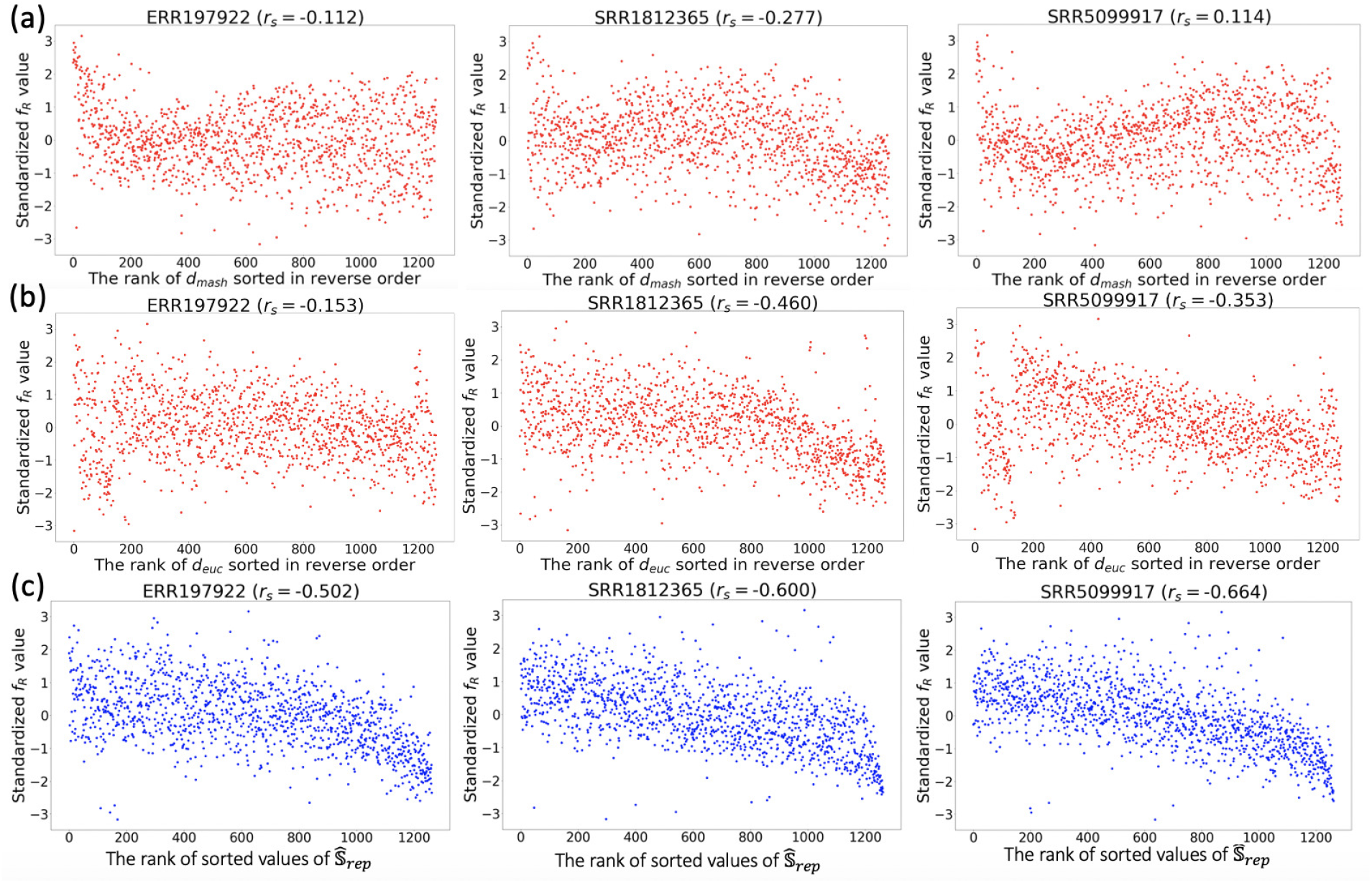
The correlation between (a) the rank of Mash similarity (negative Mash distance values −*d*_*mash*_), (b) the rank of Euclidean similarity (negative Euclidean distance values −*d*_*euc*_) or (c) the rank of sorted values of 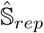 and function values 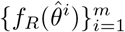 for three RNA-seq samples with accession numbers ERR197922, SRR1812365, SRR5099917. Each dot represents one representative sample (one 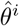). The transcript assembler 𝒜 used here is Scallop. Each y-axis is copula standardized [Salinas et al., 2020] for better visualization. *r*_*s*_: Spearman’s rank correlation coefficient.

#### 3.3.3 Rationality and performance of the data augmentation module

In this section, we first empirically validate the underlying principle of our data augmentation method: a sample *R*_*sub*_ created by subsampling reads from an original sample *R* shares a similar optimal parameter vector with *R*. We randomly select 10 samples from the representative set. For each selected sample 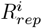, we create a subsampled version 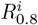 with a sampling ratio of 0.8, meaning 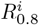 contains 80% of the reads in 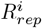 We then compute 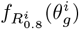 for each query point 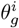 in 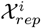 and calculate the correlation between 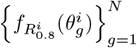 and 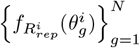, where *N* is the size of 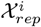 This procedure is repeated with a sampling ratio of 0.6. If the functions 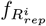 and 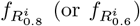 are similar, we would expect 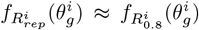 and 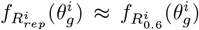 for each 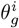 High correlations, as shown in Fig. 6, Table 7, and Table 8, indicate that for both transcript assemblers Scallop and StringTie2, when reads in 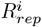 are randomly subsampled with an adequate ratio, the resulting function closely resembles the original.

**Table 7:** Pearson correlation coefficient between two sets of function values of 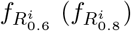 and 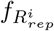 in 10 different samples. The transcript assembler used here is Scallop.

| Samples | Subsample (0.6) | Subsample (0.8) |
| --- | --- | --- |
| ERR197922 | 0.988 | 0.995 |
| SRR1812365 | 0.987 | 0.994 |
| SRR5099917 | 0.975 | 0.981 |
| SRR492054 | 0.935 | 0.975 |
| SRR346466 | 0.950 | 0.977 |
| SRR6687435 | 0.979 | 0.992 |
| SRR1782845 | 0.968 | 0.990 |
| SRR8535251 | 0.888 | 0.937 |
| SRR3932352 | 0.979 | 0.991 |
| ERR2619102 | 0.954 | 0.976 |

**Table 8:**
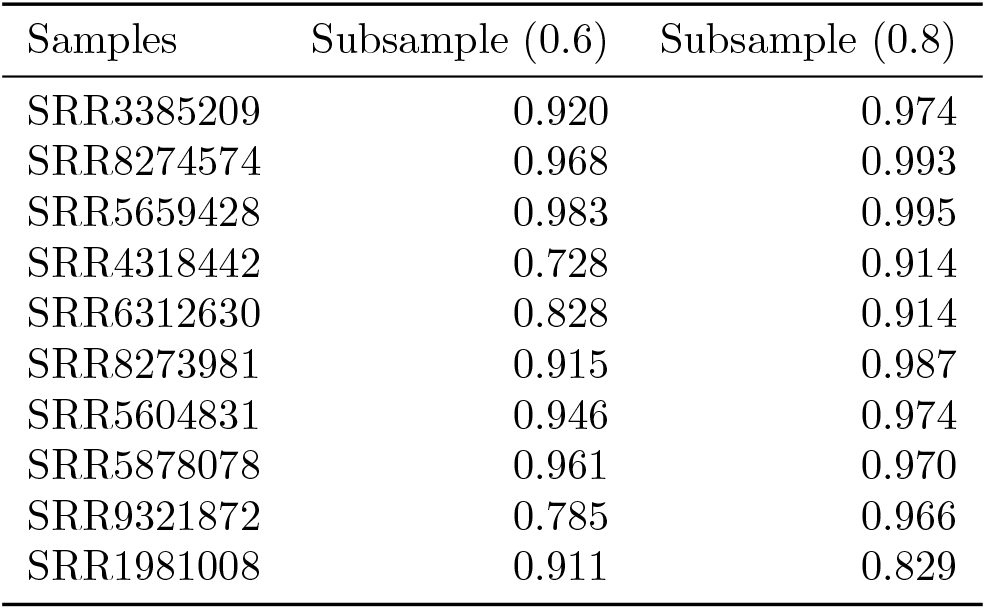
Pearson correlation coefficient between two sets of function values of 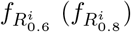 and 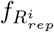 in 10 different samples. The transcript assembler used here is StringTie2.

| Samples | Subsample (0.6) | Subsample (0.8) |
| --- | --- | --- |
| SRR3385209 | 0.920 | 0.974 |
| SRR8274574 | 0.968 | 0.993 |
| SRR5659428 | 0.983 | 0.995 |
| SRR4318442 | 0.728 | 0.914 |
| SRR6312630 | 0.828 | 0.914 |
| SRR8273981 | 0.915 | 0.987 |
| SRR5604831 | 0.946 | 0.974 |
| SRR5878078 | 0.961 | 0.970 |
| SRR9321872 | 0.785 | 0.966 |
| SRR1981008 | 0.911 | 0.829 |

**Figure 6:**
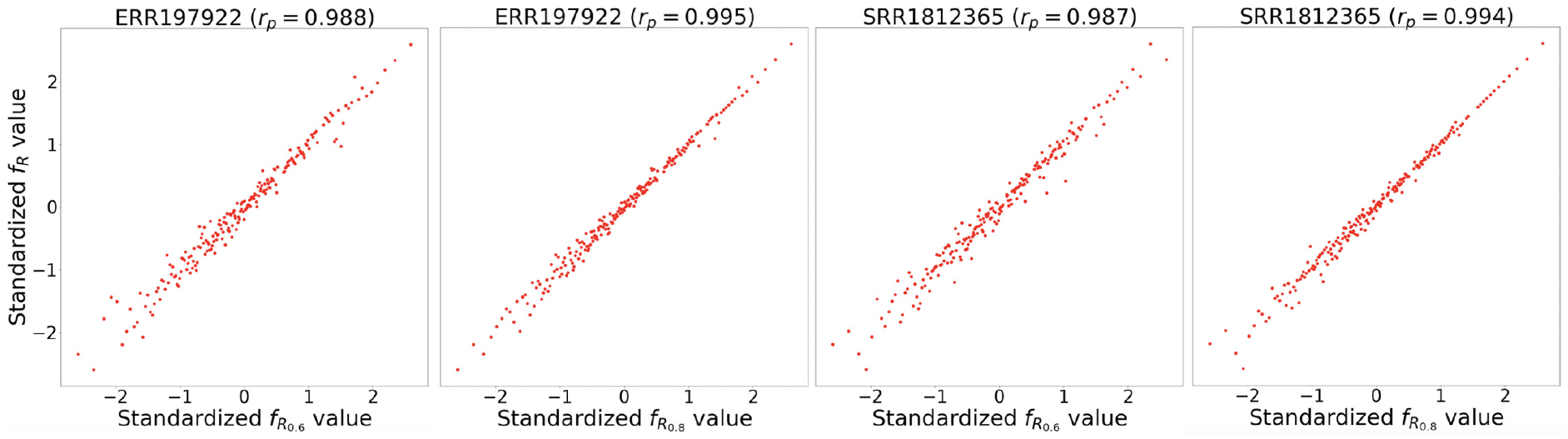
The correlation between two sets of copula standardized [Salinas et al., 2020] function values 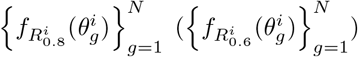 and. 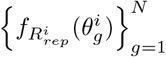 Each dot represents one parameter vector 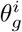. Sample accession numbers: ERR197922 and SRR1812365. The transcript assembler 𝒜 used here is Scallop. *R*_0.6_: sample from the subsampling with ratio 0.6. *R*_0.8_: sample from the subsampling with ratio 0.8. *r*_*p*_: Pearson correlation coefficient.

To confirm that the set representation of 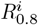 or 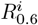 differs from 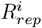 (ensuring actual data augmentation rather than duplication), we also compute the Mash distance between these two set representations. The non-zero distances reported in Table 9 and Table 10 confirm that these samples have distinct set representations. Moreover, the values in both tables are considerably lower than the average Mash distances among samples in the representative set (0.0728 for Scallop, and 0.0624 for StringTie2). These results show that the set representation of 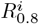 or 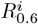 is distinct from, but similar to, 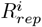, validating the efficacy of read subsampling as a reasonable data augmentation approach.

**Table 9:** The Mash distance between 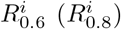 and 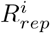 in 10 different samples. The samples here correspond to those listed in Table 7.

| Samples | Subsample (0.6) | Subsample (0.8) |
| --- | --- | --- |
| ERR197922 | 0.0080 | 0.0033 |
| SRR1812365 | 0.0083 | 0.0034 |
| SRR5099917 | 0.0071 | 0.0031 |
| SRR492054 | 0.0218 | 0.0099 |
| SRR346466 | 0.0111 | 0.0040 |
| SRR6687435 | 0.0087 | 0.0033 |
| SRR1782845 | 0.0145 | 0.0055 |
| SRR8535251 | 0.0198 | 0.0085 |
| SRR3932352 | 0.0048 | 0.0021 |
| ERR2619102 | 0.0072 | 0.0031 |

**Table 10:**
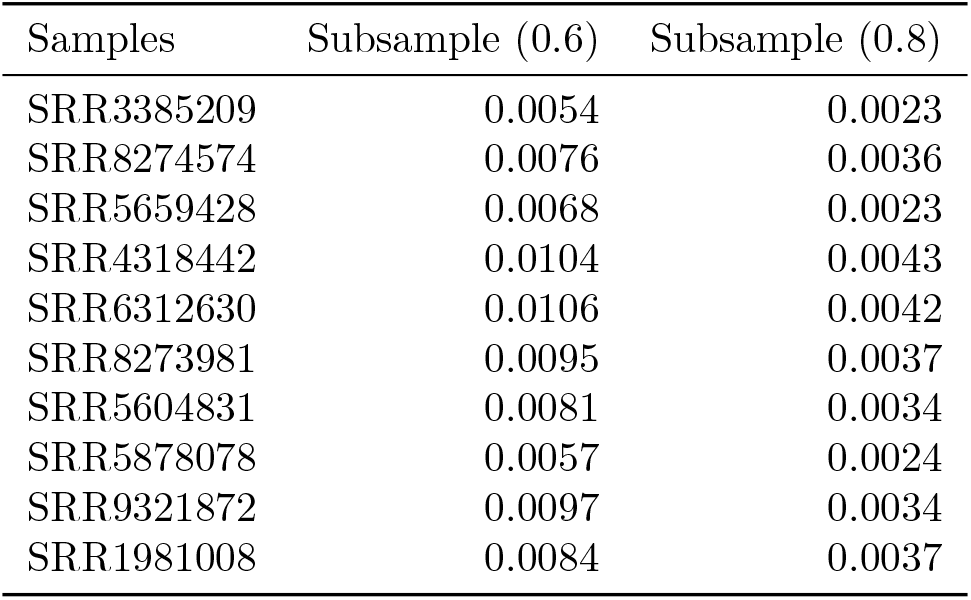
The Mash distance between 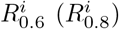 and 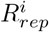 in another 10 different samples. Different from samples shown in Table 9, the samples here correspond to those listed in Table 8, which are randomly selected samples from the representative set for StringTie2.

We then empirically examine the differences in the performance of AutoTuneX with and without the implementation of data augmentation. We randomly select 150 samples from ℛ_*rep*_ to form a validation set, while the remaining samples are used for training. For each sample *R* in the validation set, we compute the Pearson correlation between the predicted similarity values and the actual predefined similarity values 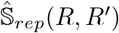, where *R*^*′*^ represents any sample in the training set. We then compare the average of these correlations across all samples in the validation set, contrasting the results obtained when training with the data augmentation module against those from training without it. Figure 7 shows the average Pearson correlations of all samples in the validation set across various training iterations. It is evident that the inclusion of data augmentation leads to significantly higher Pearson correlation values compared to the scenarios where no data augmentation is used. This trend holds true regardless of the transcript assembler employed. These findings indicate the beneficial impact and importance of the data augmentation module in enhancing the performance of AutoTuneX.

**Figure 7:**
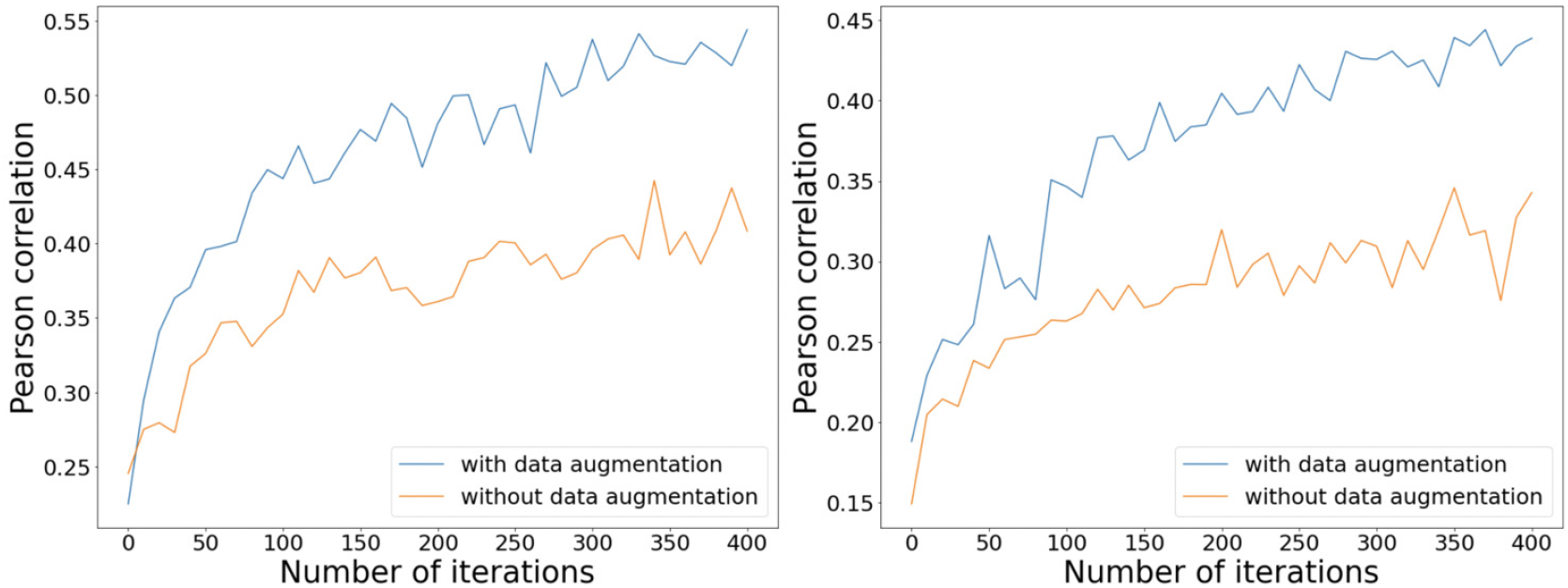
The average Pearson correlations of all samples in the validation set across various training iterations. The transcript assembler used for the left figure is Scallop, while the transcript assembler used for the right figure is StringTie2.

**Figure 8:**
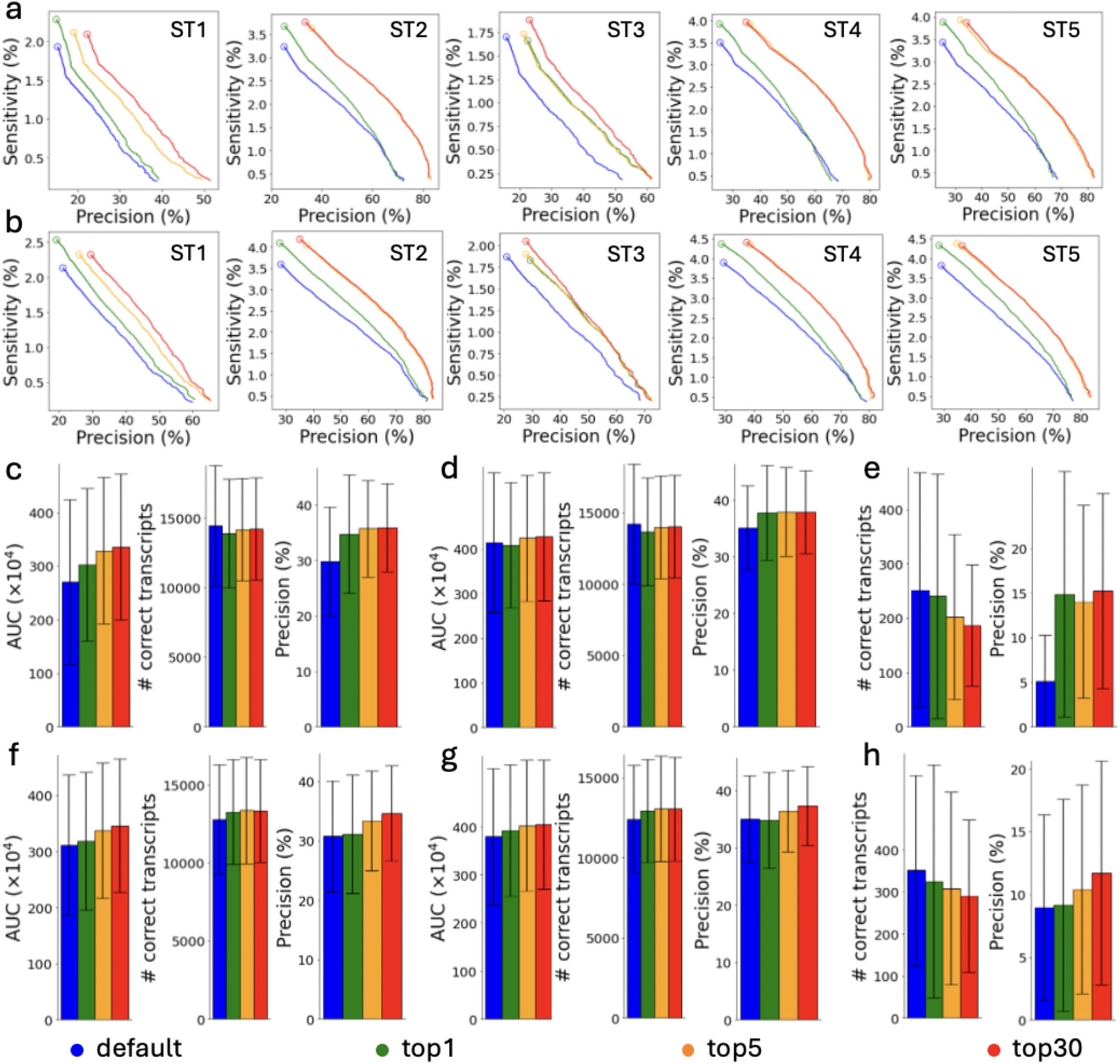
Comparisons between AutoTuneX against the default parameter settings in the top-1, top-5, and top-30 cases for ENCODE65. (c-e) are results for Scallop, and (a,b,f-h) are results for StringTie2. (a) The precision-sensitivity curves for total transcripts. The five samples are the samples that achieve the most significant AUC improvement with AutoTuneX for StringTie2. The circles represent the precision-sensitivity points with minimum coverage threshold set to 0. The accession numbers of these samples are shown in Table 11. (b) The precision-sensitivity curves for multi-exon transcripts. (c,f) The average AUC, sensitivity and precision of total transcripts running with the default minimum coverage threshold. (d,g) The average AUC, sensitivity and precision of multi-exon transcripts running with the default minimum coverage threshold. (e,h) The average sensitivity and precision of single-exon transcripts running with the default minimum coverage threshold.

**Table 11:**
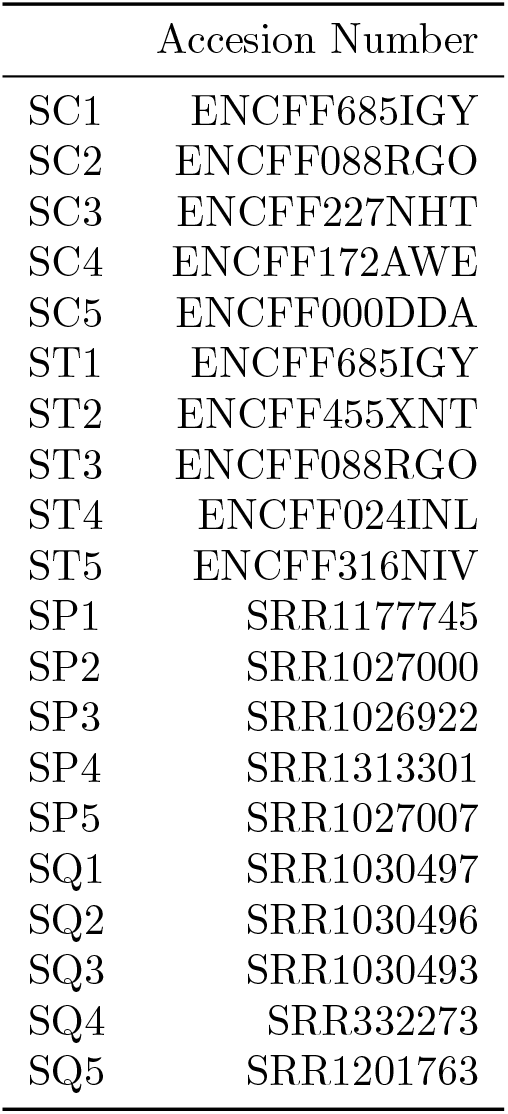
Accesion Numbers of samples shown in Fig. 1a,b and Fig. 8a,b.

**Figure 9:**
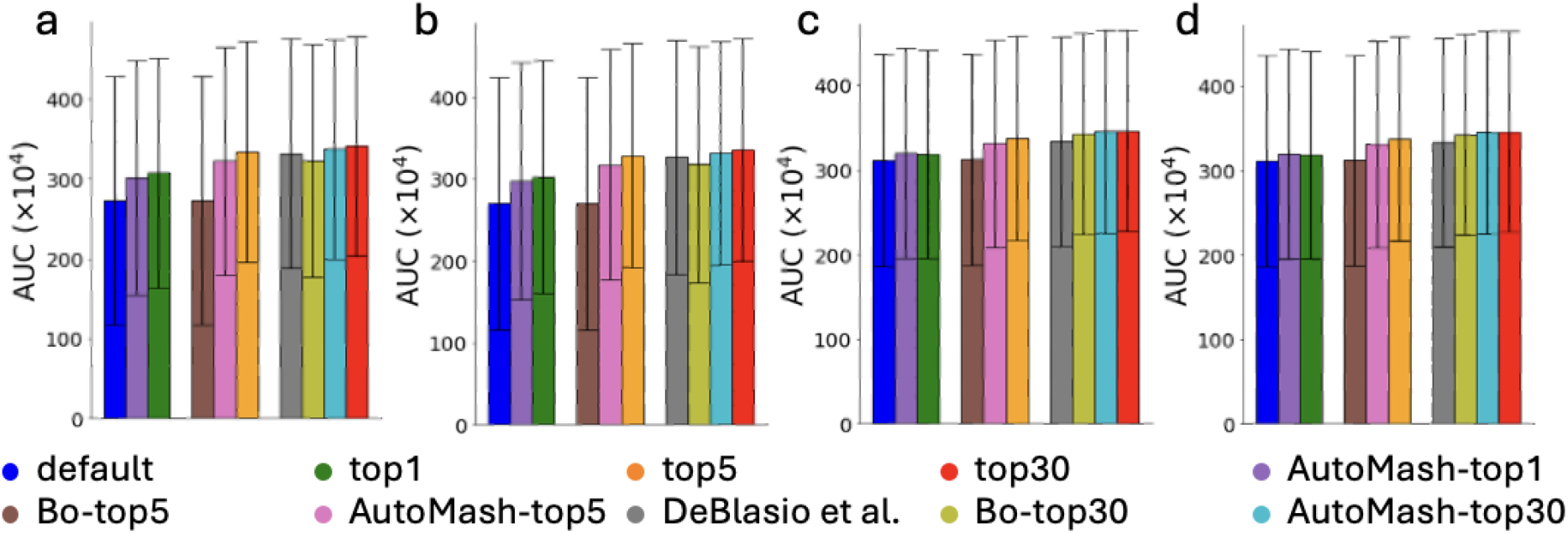
Comparisons between AutoTuneX against the method in DeBlasio et al. [2020] and two simpler heuristic methods (BO and AutoMash) in the top-1, top-5, and top-30 cases for ENCODE65. (a,b) are results for Scallop, and (c,d) are results for StringTie2. (a,c) The average AUC of total transcripts running with minimum coverage set to 0. (b,d) The average AUC of total transcripts running with the default minimum coverage threshold.

**Figure 10:**
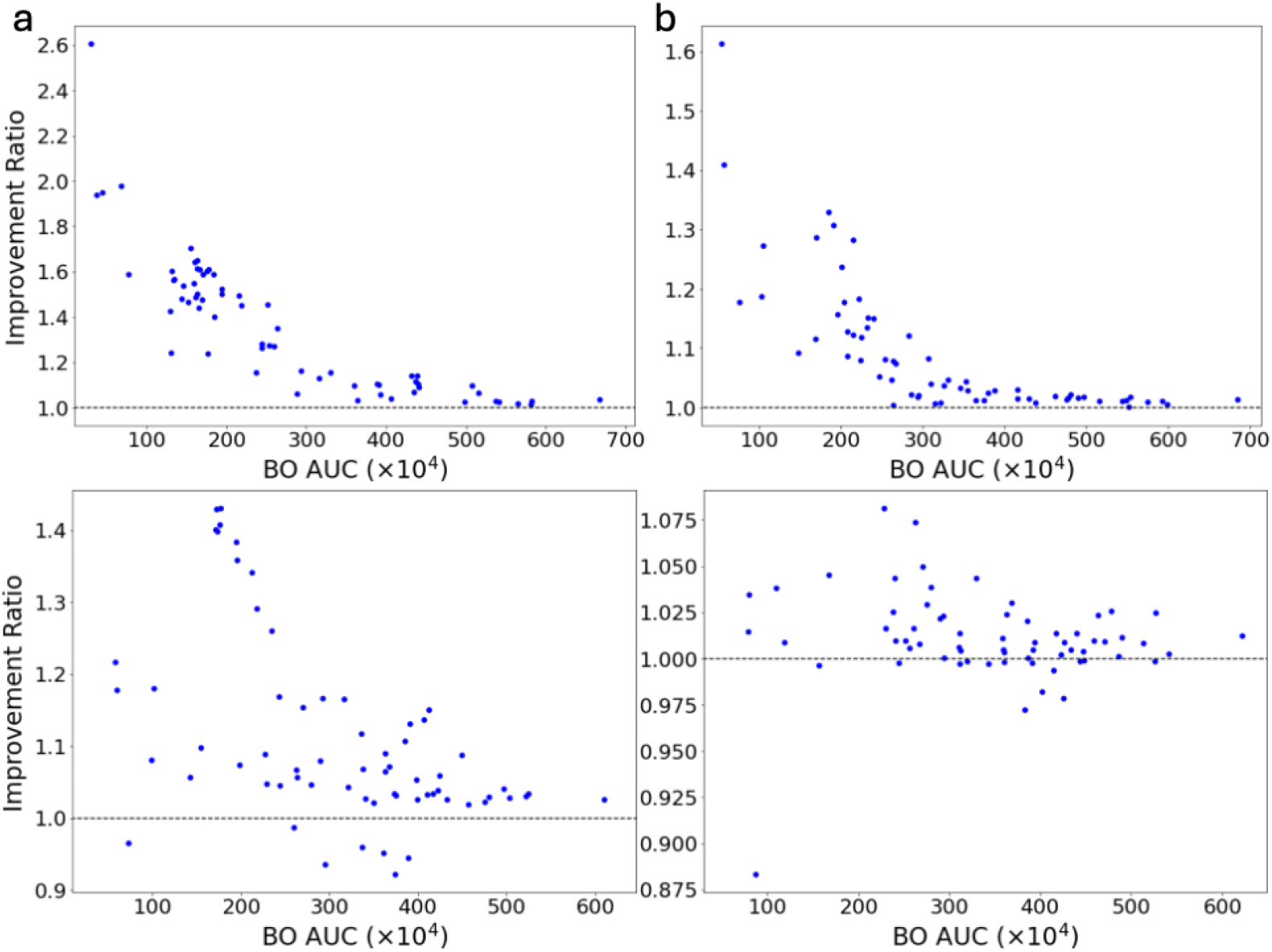
(a,b,c) Detailed comparisons between AutoTuneX against direct application of our BO framework CAWarm-BO in the top-5(b), and top-30(c) cases for ENCODE65. The transcript assembler used for the first row is Scallop and StringTie2 for the second row. Each point is one RNA-seq sample positioned by the best AUC achieved by CAWarm-BO and the ratio of the best AUC achieved by AutoTuneX relative to CAWarm-BO. A value above 1.0 indicates an improvement. We do not compare AutoTuneX against CAWarm-BO in the top-1 case because CAWarm-BO’s coordinate ascent begins from the default parameter settings, making its top-1 case identical to the default parameter settings.

**Figure 11:**
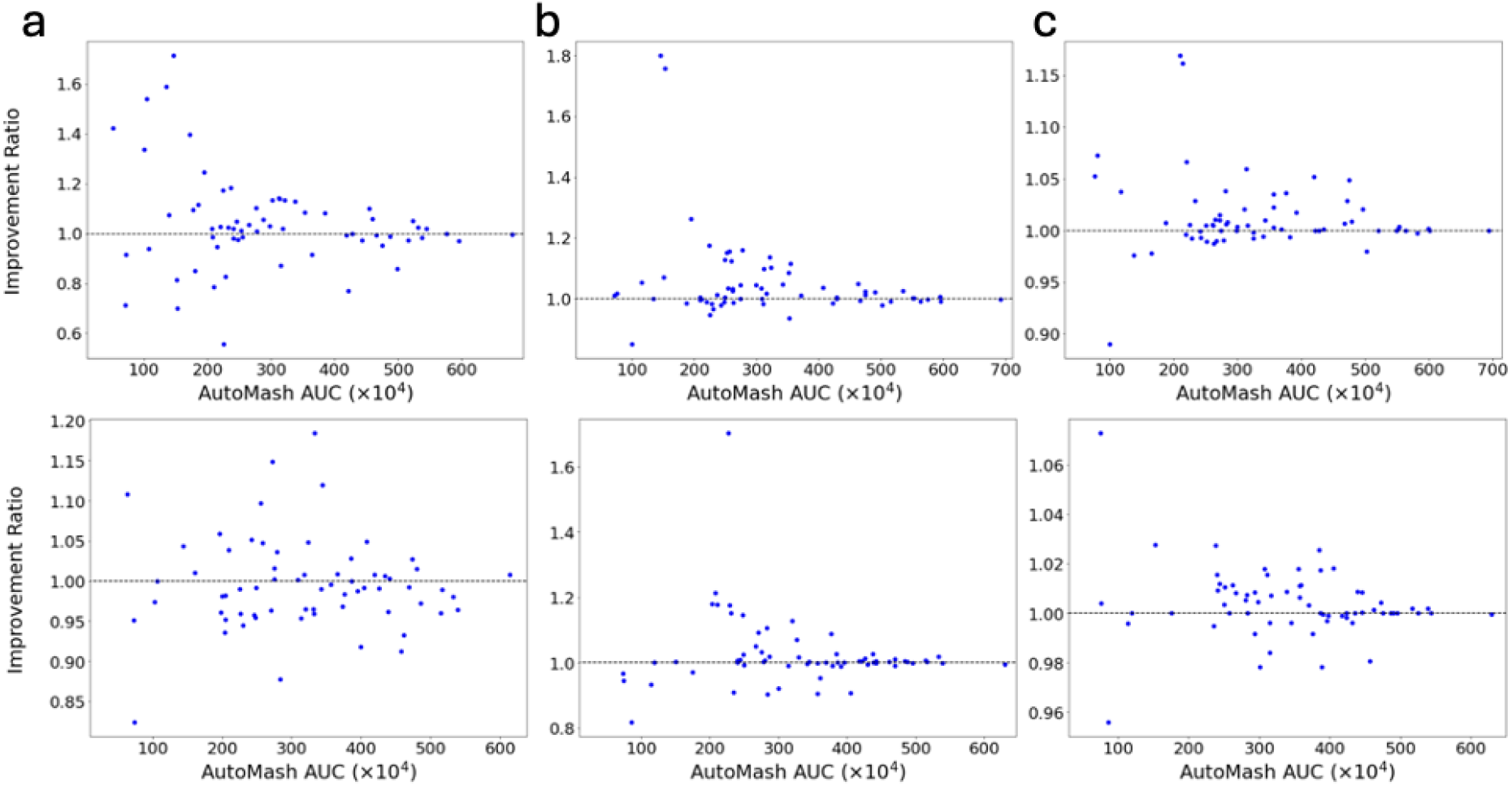
(a,b,c) Detailed comparisons between AutoTuneX against AutoMash in the top-1(a), top-5(b), and top-30(c) cases for ENCODE65. The transcript assembler used for the first row is Scallop and StringTie2 for the second row. Each point is one RNA-seq sample positioned by the best AUC achieved by AutoMash and the ratio of the best AUC achieved by AutoTuneX relative to Auto-Mash. A value above 1.0 indicates an improvement.

**Figure 12:**
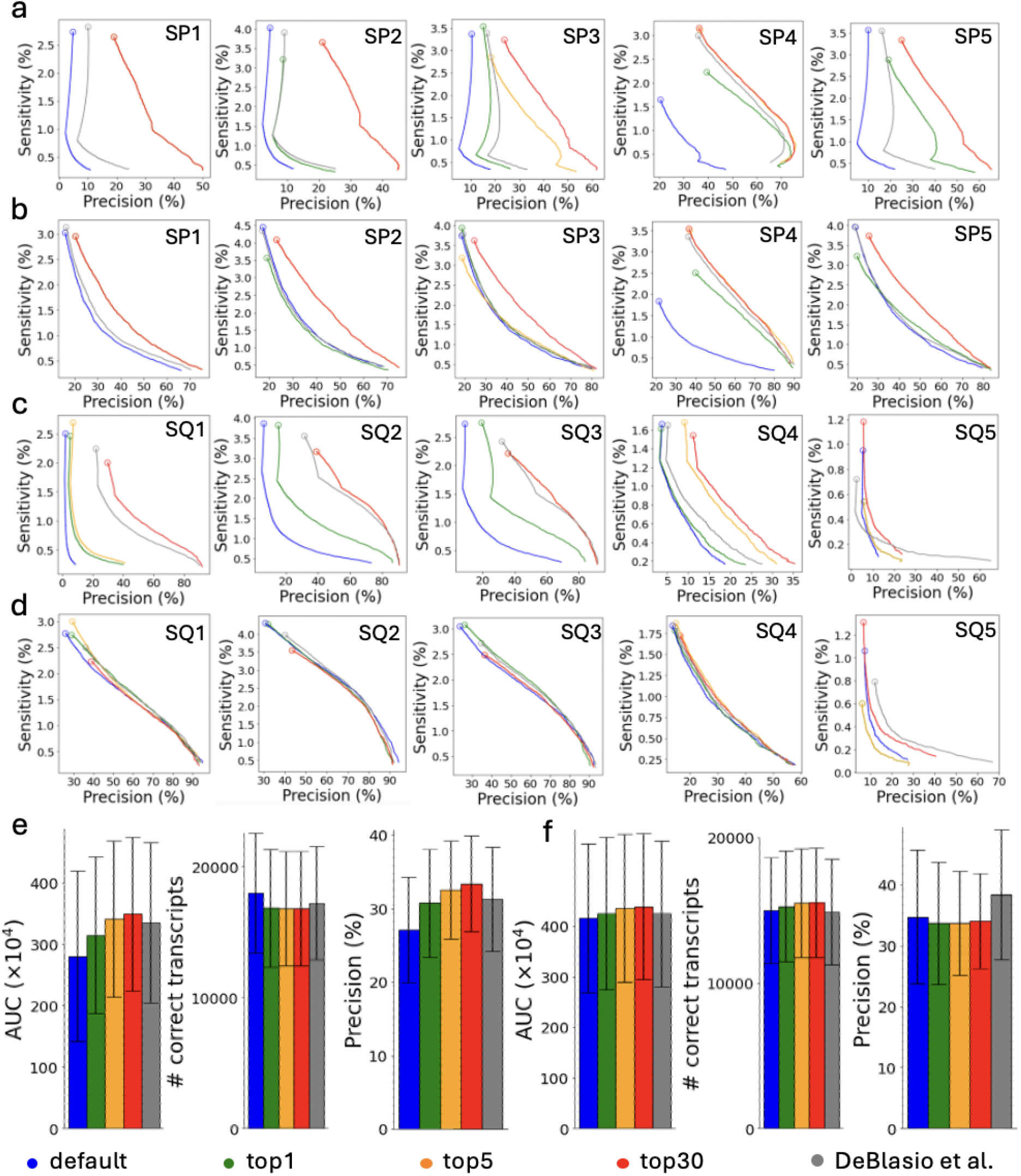
Comparisons between AutoTuneX against the default parameter settings in the top-1, top-5, and top-30 cases for SRA-test. (a,b,e) are results for Scallop, and (c,d,f) are results for StringTie2. (a) The precision-sensitivity curves for total (single- and multi-exon) transcripts. The five samples are the samples that achieve the most significant AUC improvement with AutoTuneX for Scallop. The circles represent the precision-sensitivity points with minimum coverage threshold set to 0. The accession numbers of these samples are shown in Table 11. (b) The precision-sensitivity curves for multi-exon transcripts The five samples are the same samples in (a). (c) The precision-sensitivity curves for total (single- and multi-exon) transcripts. The five samples are the samples that achieve the most significant AUC improvement with AutoTuneX for StringTie2. The circles represent the precision-sensitivity points with minimum coverage threshold set to 0. The accession numbers of these samples are shown in Table 11. (d) The precision-sensitivity curves for multi-exon transcripts The five samples are the same samples in (c). (e) The average AUC, sensitivity and precision of total transcripts running with minimum coverage set to 0 For Scallop. The error bars show the s.d. (the same for other panels). (f) The average AUC, sensitivity and precision of total transcripts running with minimum coverage set to 0 For StringTie2.

## Acknowledgements

The authors would like to thank the members of the Kingsford Group for their helpful comments throughout this project, in particular Minh Hoang, Guillaume Marçais, Laura Tung, Dan DeBlasio, and Yutong Qiu. This work was supported in part by the US National Science Foundation [DBI-1937540, III-2232121], the US National Institutes of Health [R01HG012470] and by the generosity of Eric and Wendy Schmidt by recommendation of the Schmidt Futures program. Conflict of Interest: C.K. is a co-founder of Ocean Genomics, Inc.

## Notes

### Competing Interest Statement

Carl Kingsford is a co-founder of Ocean Genomics, Inc.

### Summary of Updates

Experimental results updated, new results reported in the manuscript.

## References

William H Majoros and Steven L Salzberg. An empirical analysis of training protocols for probabilistic gene finders. BMC Bioinformatics, 5:1–12, 2004.

Martin C Frith, Michiaki Hamada, and Paul Horton. Parameters for accurate genome alignment. BMC Bioinformatics, 11:1–14, 2010.

Dan DeBlasio and John Kececioglu. Parameter advising for multiple sequence alignment. BMC Bioinformatics, 16(Suppl 2):A3, 2015a.

Sam Kovaka, Aleksey V Zimin, Geo M Pertea, Roham Razaghi, Steven L Salzberg, and Mihaela Pertea. Transcriptome assembly from long-read RNA-seq alignments with StringTie2. Genome Biology, 20(1): 278, 2019.

Dan DeBlasio, Kwanho Kim, and Carl Kingsford. More accurate transcript assembly via parameter advising. Journal of Computational Biology, 27(8):1181–1189, 2020.

Cole Trapnell, Brian A Williams, Geo Pertea, Ali Mortazavi, Gordon Kwan, Marijke J Van Baren, Steven L Salzberg, Barbara J Wold, and Lior Pachter. Transcript assembly and quantification by RNA-Seq reveals unannotated transcripts and isoform switching during cell differentiation. Nature Biotechnology, 28(5):511–515, 2010.

Mitchell Guttman, Manuel Garber, Joshua Z Levin, Julie Donaghey, James Robinson, Xian Adiconis, Lin Fan, Magdalena J Koziol, Andreas Gnirke, Chad Nusbaum, et al. Ab initio reconstruction of cell type–specific transcriptomes in mouse reveals the conserved multi-exonic structure of lincRNAs. Nature Biotechnology, 28(5):503–510, 2010.

Alexandru I Tomescu, Anna Kuosmanen, Romeo Rizzi, and Veli Mäkinen. A novel min-cost flow method for estimating transcript expression with RNA-Seq. In BMC Bioinformatics, volume 14, page S15. Springer, 2013.

Mihaela Pertea, Geo M Pertea, Corina M Antonescu, Tsung-Cheng Chang, Joshua T Mendell, and Steven L Salzberg. StringTie enables improved reconstruction of a transcriptome from RNA-seq reads. Nature Biotechnology, 33(3):290–295, 2015.

Li Song, Sarven Sabunciyan, and Liliana Florea. CLASS2: accurate and efficient splice variant annotation from RNA-seq reads. Nucleic Acids Research, 44(10):e98–e98, 2016.

Juntao Liu, Ting Yu, Tao Jiang, and Guojun Li. TransComb: genome-guided transcriptome assembly via combing junctions in splicing graphs. Genome Biology, 17(1):213, 2016.

Mingfu Shao and Carl Kingsford. Accurate assembly of transcripts through phase-preserving graph decomposition. Nature Biotechnology, 35(12):1167–1169, 2017.

Laura H Tung, Mingfu Shao, and Carl Kingsford. Quantifying the benefit offered by transcript assembly with Scallop-LR on single-molecule long reads. Genome Biology, 20(1):287, 2019.

Juntao Liu, Xiangyu Liu, Xianwen Ren, and Guojun Li. scRNAss: a single-cell RNA-seq assembler via imputing dropouts and combing junctions. Bioinformatics, 35(21):4264–4271, 2019.

Shunfu Mao, Lior Pachter, David Tse, and Sreeram Kannan. RefShannon: A genome-guided transcriptome assembler using sparse flow decomposition. PloS one, 15(6):e0232946, 2020.

Ka Ming Nip, Readman Chiu, Chen Yang, Justin Chu, Hamid Mohamadi, René L Warren, and Inanc Birol. RNA-Bloom enables reference-free and reference-guided sequence assembly for single-cell transcriptomes. Genome Research, 30(8):1191–1200, 2020.

Qimin Zhang, Qian Shi, and Mingfu Shao. Accurate assembly of multi-end RNA-seq data with Scallop2. Nature Computational Science, 2(3):148–152, 2022a.

Adam Voshall and Etsuko N Moriyama. Next-generation transcriptome assembly: strategies and performance analysis. Bioinformatics in the Era of Post Genomics and Big Data, pages 15–36, 2018.

Hyunghoon Cho, Joe Davis, Xin Li, Kevin S Smith, Alexis Battle, and Stephen B Montgomery. High-resolution transcriptome analysis with long-read RNA sequencing. PloS one, 9(9):e108095, 2014.

Guy Cochrane, Ilene Karsch-Mizrachi, Toshihisa Takagi, and International Nucleotide Sequence Database Collaboration. The international nucleotide sequence database collaboration. Nucleic Acids Research, 44(D1):D48–D50, 2016.

Brian D Ondov, Todd J Treangen, Páll Melsted, Adam B Mallonee, Nicholas H Bergman, Sergey Koren, and Adam M Phillippy. Mash: fast genome and metagenome distance estimation using MinHash. Genome Biology, 17(1):132, 2016.

Laura H Tung and Carl Kingsford. Practical selection of representative sets of RNA-seq samples using a hierarchical approach. Bioinformatics, 37(Supplement 1):i334–i341, 2021.

Jacob M Schreiber, Jeffrey A Bilmes, and William Stafford Noble. apricot: Submodular selection for data summarization in Python. J. Mach. Learn. Res., 21(161):1–6, 2020.

Bobak Shahriari, Alexandre Bouchard-Côté, and Nando Freitas. Unbounded Bayesian optimization via regularization. In Artificial Intelligence and Statistics, pages 1168–1176. PMLR, 2016.

Matthias Poloczek, Jialei Wang, and Peter I Frazier. Warm starting Bayesian optimization. In 2016 Winter Simulation Conference (WSC), pages 770–781. IEEE, 2016.

Willard I Zangwill. Nonlinear programming: a unified approach, volume 52. Prentice-Hall Englewood Cliffs, NJ, 1969.

Prannay Khosla, Piotr Teterwak, Chen Wang, Aaron Sarna, Yonglong Tian, Phillip Isola, Aaron Maschinot, Ce Liu, and Dilip Krishnan. Supervised contrastive learning. Advances in Neural Information Processing Systems, 33:18661–18673, 2020.

Yaoming Wang, Yangzhou Jiang, Jin Li, Bingbing Ni, Wenrui Dai, Chenglin Li, Hongkai Xiong, and Teng Li. Contrastive regression for domain adaptation on gaze estimation. In Proceedings of the IEEE/CVF Conference on Computer Vision and Pattern Recognition, pages 19376–19385, 2022.

Andrei Z Broder. On the resemblance and containment of documents. In Proceedings. Compression and Complexity of SEQUENCES 1997 (Cat. No. 97TB100171), pages 21–29. IEEE, 1997.

Annick Lesne, Julien Riposo, Paul Roger, Axel Cournac, and Julien Mozziconacci. 3D genome reconstruction from chromosomal contacts. Nature Methods, 11(11):1141–1143, 2014.

Manzil Zaheer, Satwik Kottur, Siamak Ravanbakhsh, Barnabas Poczos, Russ R Salakhutdinov, and Alexander J Smola. Deep sets. Advances in Neural Information Processing Systems, 30, 2017.

Luis Cedillo, Hector Richart Ruiz, and Dan DeBlasio. Exploiting large datasets improves accuracy estimation for multiple sequence alignment. bioRxiv, 2022.

Dan DeBlasio and John Kececioglu. Learning parameter sets for alignment advising. In Proceedings of the 5th ACM Conference on Bioinformatics, Computational Biology, and Health Informatics, pages 230–239, 2014.

Dan DeBlasio and John Kececioglu. Learning parameter-advising sets for multiple sequence alignment. IEEE/ACM Transactions on Computational Biology and Bioinformatics, 14(5):1028–1041, 2015b.

Chris S Magnano and Anthony Gitter. Automating parameter selection to avoid implausible biological pathway models. NPJ Systems Biology and Applications, 7(1):12, 2021.

Peter I Frazier. A tutorial on Bayesian Optimization. arXiv preprint arXiv:1807.02811, 2018.

Bobak Shahriari, Kevin Swersky, Ziyu Wang, Ryan P Adams, and Nando De Freitas. Taking the human out of the loop: A review of Bayesian optimization. Proceedings of the IEEE, 104(1):148–175, 2015.

Eric Schulz, Maarten Speekenbrink, and Andreas Krause. A tutorial on Gaussian process regression: Modelling, exploring, and exploiting functions. Journal of Mathematical Psychology, 85:1–16, 2018.

Jasper Snoek, Hugo Larochelle, and Ryan P Adams. Practical Bayesian optimization of machine learning algorithms. Advances in Neural Information Processing Systems, 25, 2012.

Aaron Klein, Stefan Falkner, Simon Bartels, Philipp Hennig, and Frank Hutter. Fast Bayesian optimization of machine learning hyperparameters on large datasets. In Artificial Intelligence and Statistics, pages 528–536. PMLR, 2017.

Eric Brochu, Vlad M Cora, and Nando De Freitas. A tutorial on Bayesian optimization of expensive cost functions, with application to active user modeling and hierarchical reinforcement learning. arXiv preprint arXiv:1012.2599, 2010.

Alonso Marco, Felix Berkenkamp, Philipp Hennig, Angela P Schoellig, Andreas Krause, Stefan Schaal, and Sebastian Trimpe. Virtual vs. real: Trading off simulations and physical experiments in reinforcement learning with Bayesian optimization. In 2017 IEEE International Conference on Robotics and Automation (ICRA), pages 1557–1563. IEEE, 2017.

Aaron Wilson, Alan Fern, and Prasad Tadepalli. Using trajectory data to improve Bayesian optimization for reinforcement learning. The Journal of Machine Learning Research, 15(1):253–282, 2014.

Ryan-Rhys Griffiths and José Miguel Hernández-Lobato. Constrained Bayesian optimization for automatic chemical design using variational autoencoders. Chemical Science, 11(2):577–586, 2020.

Diana M Negoescu, Peter I Frazier, and Warren B Powell. The knowledge-gradient algorithm for sequencing experiments in drug discovery. INFORMS Journal on Computing, 23(3):346–363, 2011.

Phuc H Le-Khac, Graham Healy, and Alan F Smeaton. Contrastive representation learning: A framework and review. IEEE Access, 8:193907–193934, 2020.

Ting Chen, Simon Kornblith, Mohammad Norouzi, and Geoffrey Hinton. A simple framework for contrastive learning of visual representations. In International Conference on Machine Learning, pages 1597–1607. PMLR, 2020.

Ching-Yao Chuang, Joshua Robinson, Yen-Chen Lin, Antonio Torralba, and Stefanie Jegelka. Debiased contrastive learning. Advances in Neural Information Processing Systems, 33:8765–8775, 2020.

Pengfei Zhang, Zhengyuan Jiang, Yixuan Wang, and Yu Li. CLMB: deep contrastive learning for robust metagenomic binning. In International Conference on Research in Computational Molecular Biology, pages 326–348. Springer, 2022b.

Geo Pertea and Mihaela Pertea. GFF utilities: GffRead and GffCompare. F1000Research, 9, 2020.

Adam Frankish, Mark Diekhans, Anne-Maud Ferreira, Rory Johnson, Irwin Jungreis, Jane Loveland, Jonathan M Mudge, Cristina Sisu, James Wright, Joel Armstrong, et al. GENCODE reference annotation for the human and mouse genomes. Nucleic Acids Research, 47(D1):D766–D773, 2019.

Xingchen Wan, Vu Nguyen, Huong Ha, Binxin Ru, Cong Lu, and Michael A Osborne. Think global and act local: Bayesian optimisation over high-dimensional categorical and mixed search spaces. arXiv preprint arXiv:2102.07188, 2021.

Alexander Dobin, Carrie A Davis, Felix Schlesinger, Jorg Drenkow, Chris Zaleski, Sonali Jha, Philippe Batut, Mark Chaisson, and Thomas R Gingeras. STAR: ultrafast universal RNA-seq aligner. Bioinformatics, 29(1):15–21, 2013.

Adam Paszke, Sam Gross, Francisco Massa, Adam Lerer, James Bradbury, Gregory Chanan, Trevor Killeen, Zeming Lin, Natalia Gimelshein, Luca Antiga, et al. PyTorch: An imperative style, high-performance deep learning library. Advances in Neural Information Processing Systems, 32, 2019.

Sergey Ioffe and Christian Szegedy. Batch normalization: Accelerating deep network training by reducing internal covariate shift. In International Conference on Machine Learning, pages 448–456. PMLR, 2015.

Diederik P Kingma and Jimmy Ba. ADAM: A method for stochastic optimization. arXiv preprint arXiv:1412.6980, 2014.

The ENCODE Project Consortium. An integrated encyclopedia of DNA elements in the human genome. Nature, 489(7414):57, 2012.

David Salinas, Huibin Shen, and Valerio Perrone. A quantile-based approach for hyperparameter transfer learning. In International Conference on Machine Learning, pages 8438–8448. PMLR, 2020.

